# Leveraging a large language model to predict protein phase transition: a physical, multiscale and interpretable approach

**DOI:** 10.1101/2023.11.21.568125

**Authors:** Mor Frank, Pengyu Ni, Matthew Jensen, Mark B Gerstein

**Author notes:** Dr. Mor Frank - end-to-end modeling framework from concept to actual methodology of both the machine learning and language models. Code writing covers all the pipeline from feature engineering, and modeling approaches to analyzing modeling results as well as writing the manuscript. Dr. Pengyu Ni - interpretation of the LLM, managing the genetic analysis and reviewing the manuscript. Dr. Matthew Jensen - preprocessed the AMP-AD dataset and obtained the gene expression matrix, Prof. Mark B Gerstein - contributed as an expert in the area of biophysics-related computational models through scientific and professional discussions as well as reviewing the manuscript. No competing interests.

## Abstract

Protein phase transitions (PPTs) from the soluble state to a dense liquid phase (forming droplets via liquid-liquid phase separation) or to solid aggregates (such as amyloids) play key roles in pathological processes associated with age-related diseases such as Alzheimer’s disease. Several computational frameworks are capable of separately predicting the formation of droplets or amyloid aggregates based on protein sequences, yet none have tackled the prediction of both within a unified framework. Recently, large language models (LLMs) have exhibited great success in protein structure prediction; however, they have not yet been used for PPTs. Here, we fine-tune a LLM for predicting PPTs and demonstrate its usage in evaluating how sequence variants affect PPTs, an operation useful for protein design. In addition, we show its superior performance compared to suitable classical benchmarks. Due to the ”black-box” nature of the LLM, we also employ a classical random forest model along with biophysical features to facilitate interpretation. Finally, focusing on Alzheimer’s disease-related proteins, we demonstrate that greater aggregation is associated with reduced gene expression in AD, suggesting a natural defense mechanism.

**Significance Statement:** Protein phase transition (PPT) is a physical mechanism associated with both physiological processes and age-related diseases. We present a modeling approach for predicting the protein propensity to undergo PPT, forming droplets or amyloids, directly from its sequence. We utilize a large language model (LLM) and demonstrate how variants within the protein sequence affect PPT. Because the LLM is naturally domain-agnostic, to enhance interpretability, we compare it with a classical knowledge-based model. Furthermore, our findings suggest the possible regulation of PPT by gene expression and transcription factors, hinting at potential targets for drug development. Our approach demonstrates the usefulness of fine-tuning a LLM for downstream tasks where only small datasets are available.

---

Protein phase transitions (PPTs) play an important role in both physiology and disease. Proteins typically exist in a soluble form, but they have the capacity to undergo a phase transition into another liquid phase, which we call droplets, or into solid condensates, typically reffered to as amyloids. Numerous studies indicate that droplets are involved in various cellular processes, such as transcription regulation, post-synaptic signaling, and genome organization (1–3). In addition to the physiological importance of droplets, emerging evidence indicates that, in neurodegenerative diseases, proteins can induce a liquid-to-solid transition to form amyloid aggregates (4, 5). Unlike droplets, which can largely revert to a soluble state, the liquid-to-solid transition is often irreversible (6).

Although it is well established that external conditions affect protein misfolding and PPT (7), understanding the fundamental factors inherently encoded by the protein sequence that drive PPT has become an additional objective. In this regard, recent studies have shown the association between droplet formation and the biophysical properties encoded by the amino acid sequence, utilizing machine learning algorithms and language models (8–11), and demonstrated that at least 40% of human proteins, known as ”droplet-drivers,” undergo spontaneous liquid-liquid phase separation (LLPS) under physiological conditions (12). However, previous modeling methods predominantly concentrate on predicting LLPS or aggregation as distinct objectives rather than describing a unified process of PPT from the soluble native state to droplet and amyloid formation. In addition, the usage of state-of-the-art techniques, such as attention-based language models, has not been fully leveraged to predict the physical mechanisms underlying diseases.

Under cellular conditions, droplets are meta-stable for most proteins, as they are kinetically trapped by a free energy barrier that inhibits the transition to a solid state (e.g., amyloid aggregation) (13). Yet, it remains unclear whether the ability to maintain this liquid meta-stable state is inherently encoded in the amino acid sequence and which biophysical properties can drive amyloid formation. Various sequence-related features affecting the formation of protein condensates have been reported, such as hydrophobic and disordered interactions (14). Here, we investigate the hypothesis that PPT is encoded by the amino acid sequence and can be directly predicted from it. Previous research using a biophysical quantity called ”binding mode entropy” (BME) demonstrated that droplet-forming proteins exhibit low BME values, indicating a strong preference for disordered binding. In contrast, amyloid-forming proteins exhibit high BME values, as they can engage in both disordered and ordered interactions (15).

Extending this approach, we calculated multiple knowledge-based features directly from the amino acid sequence and incorporated them into classical machine learning (ML) models as typically applied in the field of PPT. However, current methods predict the phase separation of proteins without a clear distinction between droplets and amyloid states (8, 9, 16, 17). This remains an important open question, as the former is mostly physiological while the latter is mostly pathological. To evaluate the performance of the classical ML-based approaches, we also fine-tuned a large language model (LLM) called ESM-2 (18) to produce contextualized embedding representations of the protein sequences. Each of the models predicted the propensity of a given protein sequence to undergo a phase transition, forming droplets and amyloid aggregates in particular (Fig. 1). By employing two distinct featurization approaches—one based on biophysical-derived features and another on contextualized embedding vectors—we explored how well the LLM can learn PPT in comparison to essential features learned by classical ML models. To accomplish this, we trained the two models, in classification tasks A and B, dividing our datasets into training and test sets in a ratio of 80:20, respectively, and analyzed the learning criteria of each model.

**Fig. 1.**
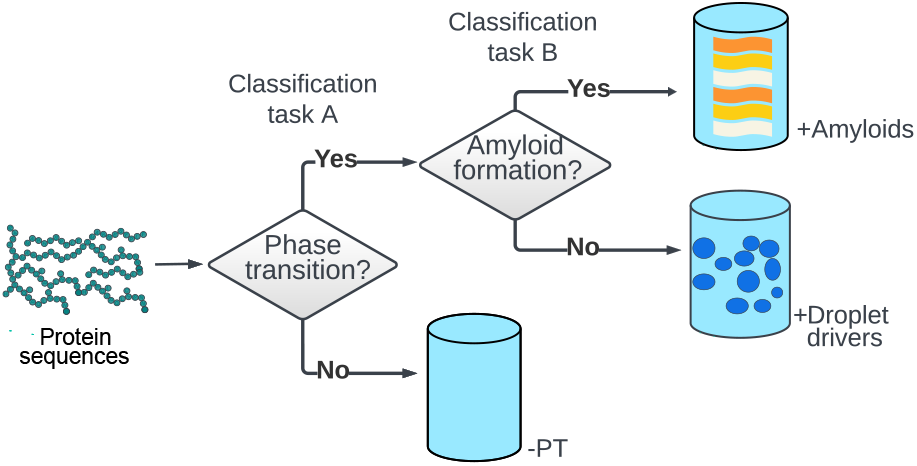
Modeling approach for PPT. In classification task A, proteins exhibiting experimental evidence of undergoing a phase transition, forming either droplets or amyloids, are consolidated into a single dataset (+Droplet drivers and +Amyloids). Phase transition propensity is predicted versus the preference to maintain the native soluble state (-PT). In classification task B, the unified dataset is utilized to predict the propensity to form droplets versus amyloid aggregates.

Next, we demonstrated the value of attention mechanism in highlighting important regions that mostly affect PPT predictions. This suggests a convenient way to guide experiments by focusing on specific regions where variants are most likely to affect PPT propensity.

Finally, in an effort to establish a link between PPT propensity and gene expression, we compared the gene expression of amyloid-beta precursor protein (APP) and tau proteins—both known to contribute pathologically to Alzheimer’s disease (AD)—to better understand how PPT is regulated under pathological conditions. Our results suggest the possibility that the tendency to form amyloid aggregates, as well as liquid droplets, is encoded by the amino acid sequence. Furthermore, this process appears to be self-regulated by gene expression, thereby controlling protein aggregation, as seen in the case of AD.

## Results and Discussion

### Leveraging ESMFold, a large language model, for predicting protein phase transitions

Recently, pre-trained language models employing deep bidirectional attention have proven to be useful in embedding biological sequences (18, 19). Here, we leveraged the evolutionary scale ESMFold model, which was trained on a large database of evolutionarily diverse protein sequences (20), for predicting PPT. Unlike the AlphaFold model (21), which employs a deep neural network architecture to facilitate information exchange between protein sequences (i.e., an Evoformer block), ESMFold utilizes a masked transformer to analyze a single sequence. This distinction makes ESMFold an order of magnitude faster than AlphaFold in predicting protein structures. Leveraging this advantage, we adopted ESMFold for predicting PPT in the above-mentioned classification tasks. Specifically, we used the pre-trained ESM-2, a transformer-based language model that is part of the ESMFold architecture. Additionally, we incorporated a linear layer on top of the model to handle the classification task instead of using the folding module (Fig. 2).

**Fig. 2.**
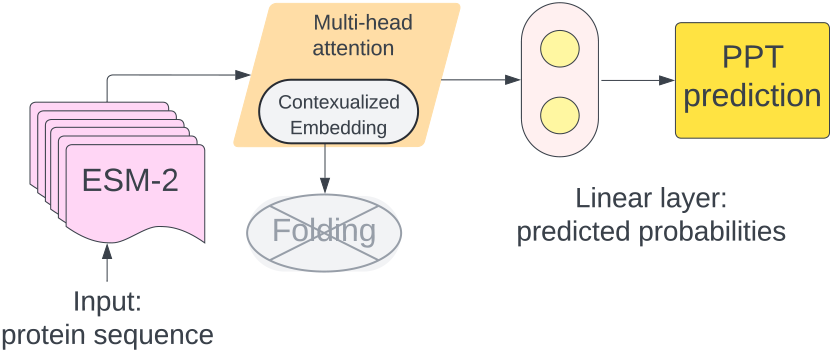
ESM-2, a pre-trained LLM via masking, is utilized to predict PPT. The folding module is removed from the original architecture of the ESMFold model, while a linear head is added on top for employing a classification task.

In the following sections, we demonstrate that this modeling approach for predicting PPT exhibits better performance than classical model benchmarks (Supplementary Tables S1 and S2). The discriminative power of the LLM versus the random forest (RF) model is demonstrated through principal component analysis (PCA) of the embedding vectors extracted from the last layer of the LLM versus the PCA of the biophysical features (Fig. 3).

**Fig. 3.**
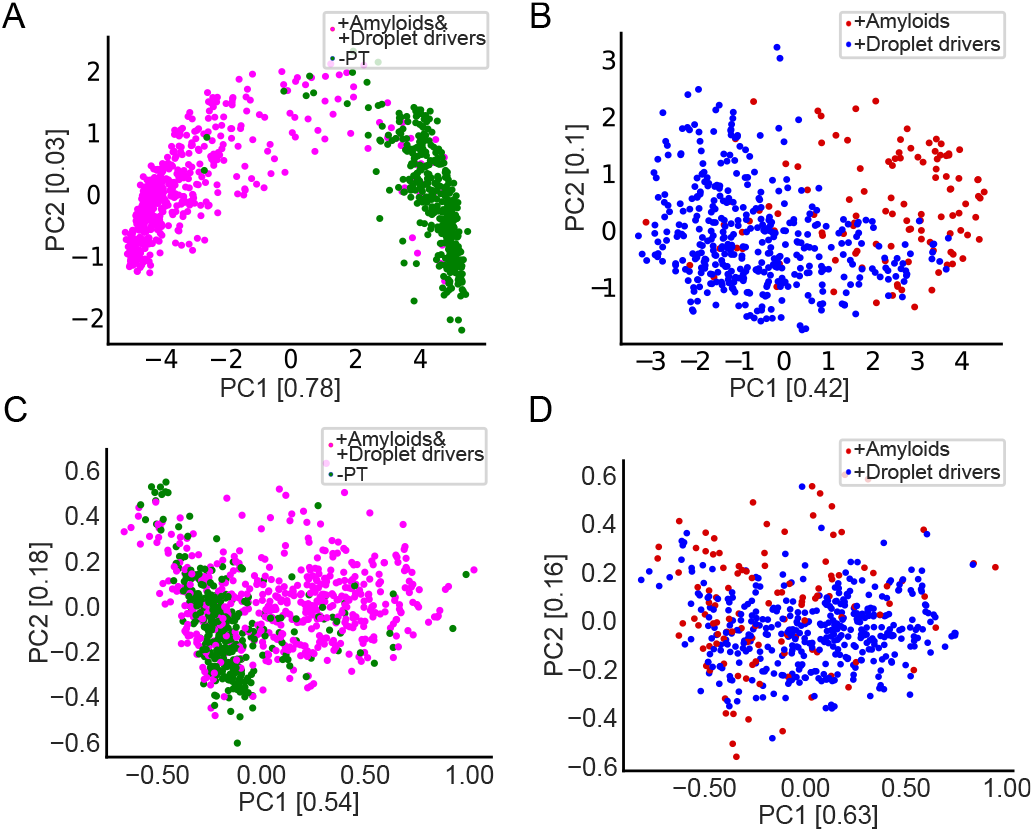
PCA of the embedding vectors, extracted from the LLM, and the biophysical features for both classification tasks. Each principle direction is presented along with its explained variance. The embedding vector of each protein sequence is the mean of all the 320-dimensional embedding vectors representing the amino acids in the sequence and was extracted from the last hidden layer. (A) and (B) The PCA of the embedding vectors is presented for classification tasks A (+Droplet drivers and +Amyloids versus -PT) and B (+Droplet drivers versus +Amyloids), correspondingly. (C) and (D) The PCA of the biophysical features of the two classification tasks, A and B, are shown.

### Database construction of the biophysical features

We used two databases containing full protein sequences that represent the propensity to form droplets (*+Droplet drivers*) and amyloid aggregates (*+Amyloids*), following the process in (15) with no further modifications. It should be noted that our analyses excluded proteins observed to exhibit both droplet and amyloid formation. This decision was driven by the limited availability of data, with only 20 proteins reported to possess such dual capabilities. However, our model captures the degree to which proteins are prone to form droplets as a prior step for amyloid formation. In addition, we restricted our analyses to droplet drivers to specifically focus on the biophysical properties of proteins that do not require an interacting partner to undergo LLPS. Moreover, we constructed a negative database of well-structured proteins randomly chosen from the database used in (8). These proteins are highly unlikely to undergo LLPS and are not reported to form amyloid aggregates, thus demonstrating a minimal propensity for undergoing a phase transition (*-PT*) (Materials and Methods).

We computed the following biophysical features, which are commonly used in the field of PPT (8, 15, 22, 23), considering multiple protein scales (Fig. 4 (A-E), and Table 1) and kept a limited set of the most significant ones (SI, Appendix, Fig. S12). First, we identified disordered residues (Materials and Methods) using the iupred3 algorithm (24). In accordance with (15), our analyses revealed that +Droplet drivers had a higher fraction of disordered regions compared to +Amyloids, which preserved a more ordered structure as demonstrated in Fig. 4A. Second, to account for the 3D structure of the proteins, we used the Protein Data Bank (PDB) files extracted from the AlphaFold (29) database or used the ESMFold model (20) and calculated the gyration radius of each protein sequence (Materials and Methods). We concluded that +Amyloids tend to have a more compact structure, consistent with their tendency to have more ordered interactions compared to +Droplet driver proteins, which primarily demonstrate disordered interactions (Fig. 4B). This compact structure maximizes interactions potentially encoded in the amino acid sequence and promotes aggregation. Third, we calculated the instability index, *II*, of a given protein sequence as shown in Fig. 4C, (Materials and Methods). +Droplet drivers are associated with high *II* in accordance with their unstable disordered structure under physiological conditions (30), as opposed to -PT, which shows high stability. The +Amyloids showed a relatively low *II*, aligning with their propensity to undergo misfolding while maintaining a stable structure of beta sheet-rich assemblies (31). This is in contrast to +Droplet drivers, which are associated with a high fraction of disordered regions and a lack of a stable 3D structure (32). Next, we estimated the probability of solubility for all protein sequences using the solubility weighted index, *SWI* (Materials and Methods). We found that +Droplet drivers and -PT exhibited a slightly higher probability of solubility than +Amyloids (Fig. 4D). This characteristic could be linked to their important cellular-level functions while evading aggregation. Finally, we calculated the mean hydrophobicity using the Kyte and Doolittle hydropathy scale for each protein sequence (Fig. 4E). We found that +Droplet drivers showed the lowest values, suggesting an essential role of hydrophilic interactions in the LLPS process. These interactions allow the chains to remain disordered, maintaining the condensates in a liquid state rather than a solid one (14). In contrast, +Amyloids exhibited higher values associated with hydrophobic interactions, which promote aggregation.

**Table 1.** Biophysical features.

| Feature | Reference |
| --- | --- |
| 1. Fraction of disordered regions | (24) |
| 2. Gyration radius | (25) |
| 3. Instability index | (26) |
| 4. Probability of solubility | (27) |
| 5. Avg. hydrophobicity | (28) |

**Fig. 4.**
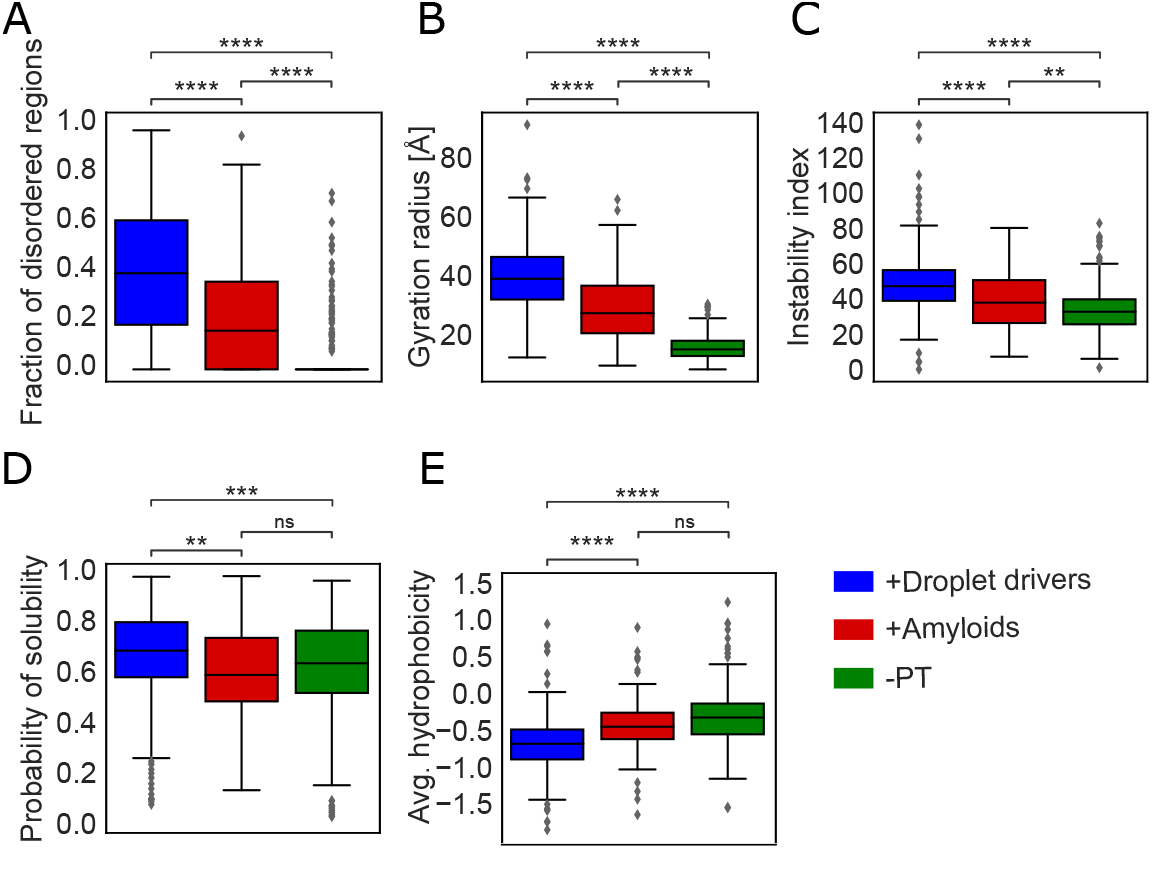
Wide range of biophysical features demonstrating the differences between the three datasets. (A) Fraction of disordered regions, (B) Gyration radius, (C) Instability index, (D) Probability of solubility, and (E) Average Hydrophobicity. A significant difference was found between +Droplet drivers and +Amyloids. When comparing -PT to +Droplet drivers or +Amyloids, a significant difference emerged in most cases. As expected, +Droplet drivers show a more disordered structure compared to +Amyloids, while -PT shows almost no disordered regions (33). The gyration radius of +Amyloids is lower compared to +Droplet drivers. +Amyloids show reduced instability index values compared to +Droplet drivers. Overall, the solubility probability of +Droplet drivers and -PT is higher compared to +Amyloids. +Droplet drivers show the lowest average hydrophobicity, while -PT shows the highest. Significance was tested using the Welch statistical t-test. **** p <= 1.00e-04, *** 1.00e-04 < p <= 1.00e-03, ** 1.00e-03 < p <= 1.00e-02, and ns is no significance.

### Prediction of phase transition

We first predicted whether a given protein would undergo a phase transition or prefer to remain in the native state. For this, we considered the +Droplet drivers and +Amyloids as a unified dataset, comparing them against the -PT dataset (classification task A). If a protein is predicted to undergo PPT, it is further evaluated to determine whether it is more likely to form +Droplet drivers or +Amyloids (classification task B).

For both classification tasks, two clusters could be distinguished through PCA performed on either the embedding vectors or the biophysical features. Remarkably, for both classification tasks, a clearer distinction between the two classes was achieved when using the LLM embedding vectors (Figs. 3A and 3B). This result emphasizes the importance of latent relationships derived from the protein sequence, which affect the likelihood of PPT occurrence. However, it is evident that more learning criteria are still needed to distinguish between +Droplet drivers and +Amyloids, as shown in SI Appendix, Fig. S11.

A notable separation was achieved between the unified dataset of +Droplet drivers and +Amyloids versus -PT (Figs. 3A and 3C). Additionally, we observed a relatively clear separation between +Droplet drivers and +Amyloids (Figs. 3B and 3D). This difference supports the idea that the trade-off between the proper functionality of droplets and the risk of forming solid aggregates, i.e., amyloid, is governed by the biophysical properties encoded in the amino acid sequence. Nonetheless, distinguishing between +Droplet drivers and +Amyloids was more challenging, as droplet-promoting proteins can transition into an amyloid state under certain conditions, as demonstrated in a recent study of LLPS-prone proteins (34).

To evaluate the performance of the two models, we performed five-fold cross-validation in a stratified manner. We used accuracy, the arithmetic mean of recall and precision, and the area under the receiver operating characteristic curve (AUROC) to evaluate the RF model and the LLM, as shown in Fig. 5. The two models successfully distinguished between proteins that have a high propensity to undergo a phase transition, whether droplets or amyloid aggregates, as opposed to those that tend to remain in their native state (Fig. 5A). The distinction between +Droplet drivers and +Amyloids was also fairly notable (Fig. 5B). This classification task is more challenging, as +Amyloids can have both disordered and ordered binding modes (i.e., interactions) associated with both liquid and solid condensates. Remarkably, all metric quantities were higher in the LLM than in the RF model for both classification tasks. The high predictive power of the LLM is in line with the PCA presented in Figs. 3A and 3B, where a clearer distinction between the two classes was achieved by the embedding vectors than by the biophysical features. This indicates that the LLM successfully identifies regions that drive PPT, suggesting that various biophysical properties, beyond those presented in Fig. 4, are encoded by the amino acid sequence.

**Fig. 5.**
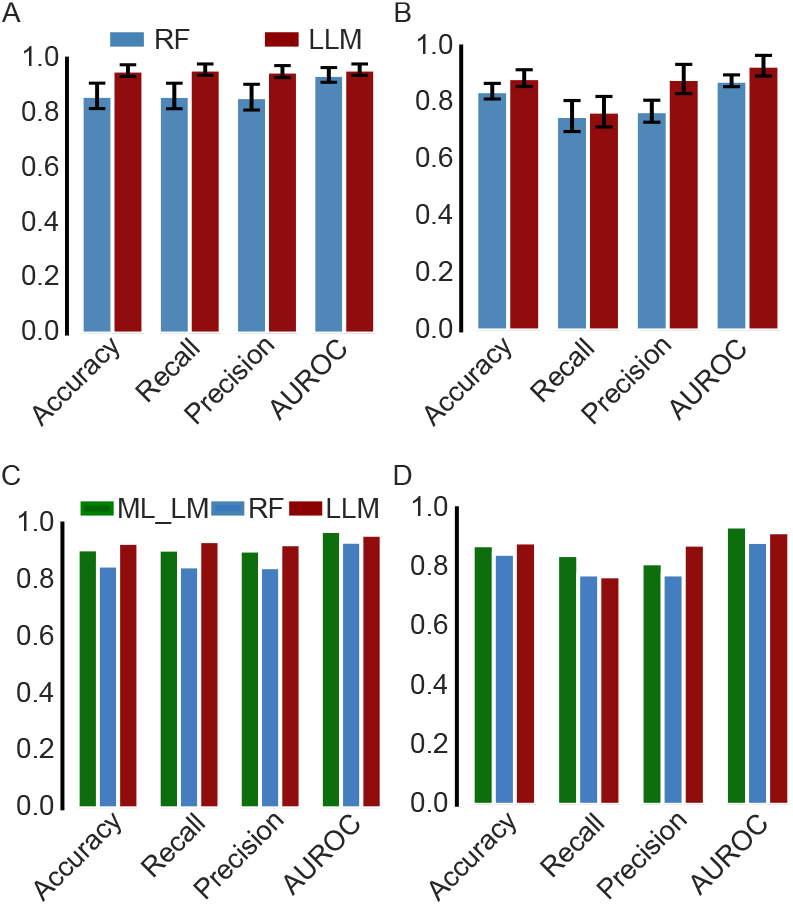
Metrics evaluating the RF model and the LLM. (A) and (B) Five-fold cross-validation showing the mean and standard deviation. (C) and (D) Performance on an external test set. (A) Classification task A: +Droplet drivers and +Amyloids versus -PT. Both models showed consistently high performance for all four metrics, which were all above 0.8 for RF and 0.9 for the LLM. (B) Classification task B: +Droplet drivers versus +Amyloids. Performances were still high, although lower than classification task A due to the challenging distinction between the two classes. (C) The performance of classification task A, +Droplet drivers and +Amyloids vs. -PT, showed consistently high measures, all above 0.9 for both the LLM and the ML_LM. (D) For classification task B, +Droplet drivers vs. +Amyloids, all four metrics were higher than 0.8 for the ML_LM.

Next, we created an ensemble of the RF and the LLM models (ML_LM model) by averaging their predicted probabilities (Materials and Methods). To evaluate the performance of the ML_LM model, we calculated the same metrics for the test set. In both classification tasks, the LLM demonstrated better performance than the RF model in almost all metrics, while the ML_LM model achieved the highest AUROC (Figs. 5C and 5D). Furthermore, the ML_LM model significantly improved the ability to identify protein sequences belonging to either +Droplet drivers or +Amyloids (Fig. 5D, Recall metric).

To gain a deeper understanding of how the LLM makes PPT predictions, we extracted self-attention maps for tau and APP (Fig. 6). In both classification tasks, the LLM identified patterns within the protein sequences, indicating that certain combinations of amino acids exhibit greater significance than others. Specifically, for classification task A, local attention was useful in predicting PPT (Figs. 6A and 6D). However, to distinguish between droplet and amyloid-forming proteins, a broader contextualized interaction was needed. This suggests that the model seeks additional information to make predictions (Figs. 6B and 6E), likely due to the complexity of this classification task. This difference between the learning criteria of the two tasks, as revealed by the attention maps, is quantified by the locality parameter *Lw*, which is defined as the ratio between the median attention score and the mean distance of the top 5 scores from the target amino acid (located on the diagonal) within a 5×5 sliding window around the target amino acid (Eq. 6). The intuition of this score is that it increases when attention is high (numerator) or the highest scores are close to the target (denominator). Local attention is associated with high Lw values, whereas a sparse attention pattern is linked to lower values. The *Lw* line plots in Figs. 6C and 6F show how attention scores vary across amino acid positions for tau and APP. In classification task B, both proteins exhibit lower Lw values compared to task A, with a significant difference between them (p<0.05). A similar trend was observed in FUS, hnRNPA1, and G3BP1, as presented in SI Appendix, Fig. S10.

**Fig. 6.**
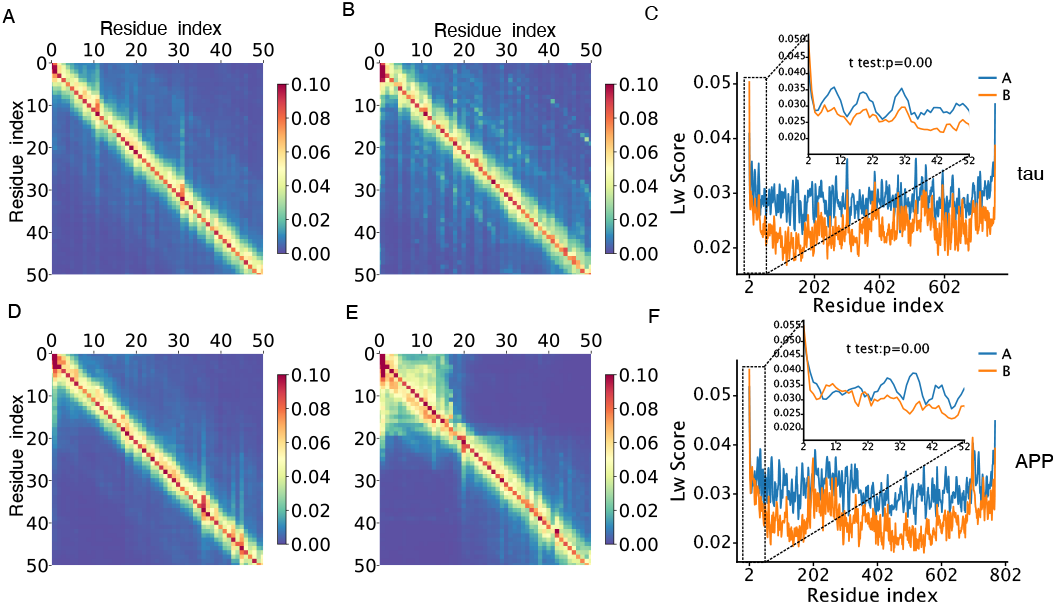
Self-attention maps for the first 50 amino acids in sequence for tau and APP proteins extracted from the LLM (left panel: classification task A; middle panel: classification task B). Attention scores referring to each amino acid (key tokens) are presented on the x-axis. The y-axis represents the target amino acids (query tokens) for which an attention score vector is computed. The attention scores were computed as the average across the 20 attention heads. These scores correspond to the level of attention that the query should have for each key. (A), (B), and (C) refer to tau. (D), (E), and (F) refer to APP. (A) and (D) exhibit strong local patterns where attention scores are close to being a band matrix with interest concentrated in small regions near the diagonal, indicating very local self-attention contextualized interactions. In (B) and (E) attention scores are more sparse, indicating broader contextualized interactions. (C) and (F) The difference between the local and sparse attention is presented by the Lw line plots.

We next calculated the contribution of each biophysical feature using the Gini score (35), as presented in Fig. 7. The most important features for both tasks are the instability index, the fraction of disordered regions, and the gyration radius. Classification task A is mostly dominated by the ”fraction of disordered regions” feature, while in classification task B, the contribution of all features is more evident. In addition, we observed that classification task A is mostly governed by structural features, while classification task B is mostly driven thermodynamically, reflected by the instability index as the most important feature. Taken together, in the RF model, the two classification tasks are dictated by different learning criteria while preserving the same important features (the three top important features). This distinction in the learning criteria of the two tasks is evident also in the LLM attention maps, where classification task A shows local attention associated with high *Lw* scores, whereas classification task B exhibits sparse attention linked to lower *Lw* scores.

**Fig. 7.**
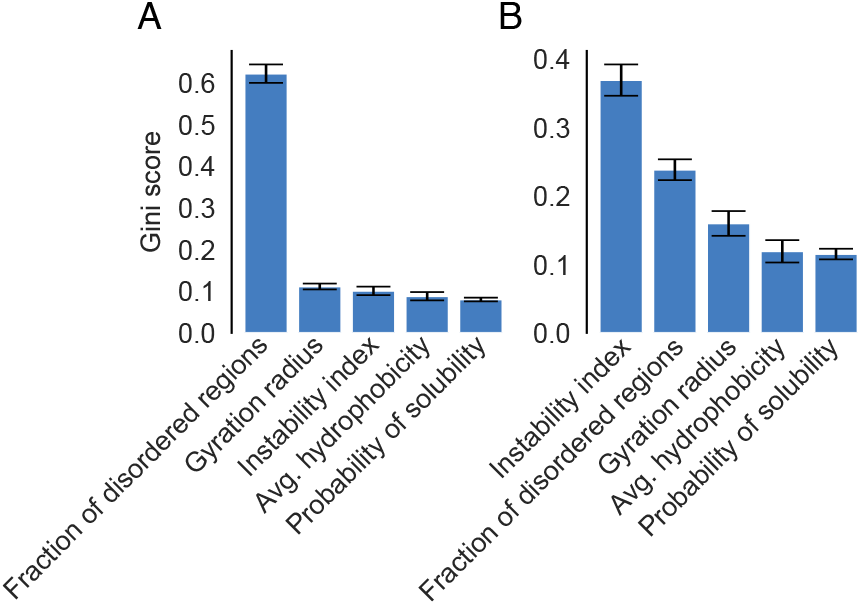
Feature importance calculated for the two classification tasks by the mean Gini score derived from the 5-fold cross-validation, along with the standard deviation. (A) For classification task A,+Droplet drivers and +Amyloids versus -PT, the fraction of disordered regions shows significant discrimination power, while other features contribute similarly. (B) In classification task B, +Droplet drivers versus +Amyloids, the instability index and the fraction of disordered regions are the most important features, followed by a gradual decrease in the Gini score of the other features. This emphasizes their accumulated contribution to the prediction.

To quantify a protein’s inherent tendency to preferentially adopt a specific state, we introduced a scoring function called the *“Transition score,”* which is defined by a log-odds ratio using the probability of undergoing PPT (Materials and Methods). First, we calculated the probability of transitioning from the native state into either droplets or amyloid states (classification task A). We applied Eq. 4 to the unified dataset of +Droplet drivers and +Amyloid versus -PT, where *p* is the probability of being in the unified +Droplet drivers and +Amyloids class. Our analysis resulted in a clear separation, with the largest densities located in negative and positive ranges, suggesting the predictability of PPT (Fig. 8A). Second, we used this measure to distinguish between +Droplet drivers and +Amyloids by using *p* in Eq. 4 as the probability of being in the +Amyloids class (classification task B). In this classification task, a lower transition score suggests that the protein exhibits more disordered interactions than ordered ones, potentially promoting droplet formation prior to amyloid formation. We achieved reasonable discrimination between these two classes (Fig. 8B), with some overlap between the two distribution scores. However, high densities were located within both negative and positive ranges.

**Fig. 8.**
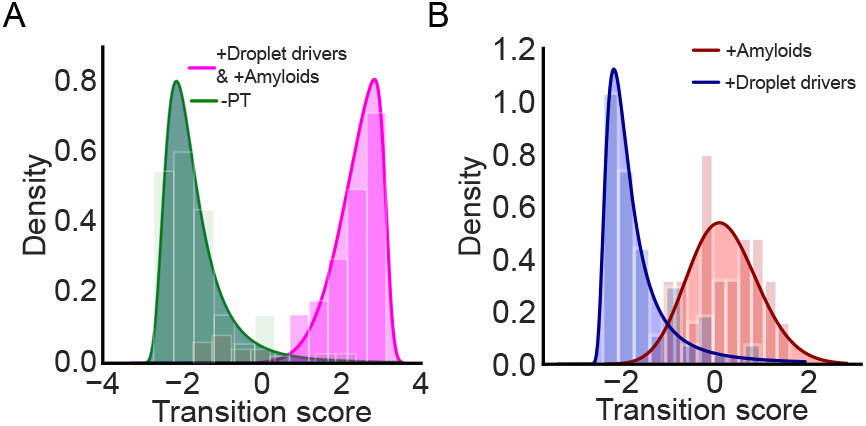
Probability density of the transition scores, along with the fitted alpha distributions (solid line), for the two test sets using ML_LM. (A) Transition scores of classification task A: +Droplet drivers and +Amyloids versus -PT. A clear distinction was achieved between the distributions of the two classes, as they are located separately in the negative and positive ranges. (B) Transition scores of classification task B: +Amyloids are associated with higher transition scores, which are mostly in the positive range.

### Towards variant design accounting for phase transition propensity

Modeling the effect of sequence variants on PPT is important for understanding the mutations that contribute to aggregation. This, in turn, is useful in various protein design applications. As an illustration, we show the results of running our LLM model, which inherently captures local residue information (8), on a number of mutations in the three proteins in SI Appendix, Figs. S5-S9. In the calculations, we can see how some mutations favor slightly droplet formation or aggregation and how the mutations are also manifest changes in the attention score. Next, we computed transition scores for mutated sequences of *Aβ*42 protein that were experimentally shown to reduce aggregation (36) using our ML_LM model (Materials and Methods). We observed that most of the variants affecting aggregation are tightly clustered within a local region within a third of the *Aβ*42 sequence (701-713 positions in APP). Interestingly, this region also has elevated attention in both classification tasks highlighting its importance for the predictions and also motivating experimental exploration of potential mutants (Figs. 9A-C). The variants in this region are all positive for classification task A (Fig. 9D). For task B, most variants tend to reduce aggregation (30/36, Fig. 9E). This is in line with the finding that lower aggregation score is associated with greater propensity for folding (as experimentally measured, Fig. 9F; dataset S7 and dataset S8). These task-B variants might sustain the droplet state longer as demonstrated in a recent study involving *Aβ*42 as discussed in (37).

**Fig. 9.**
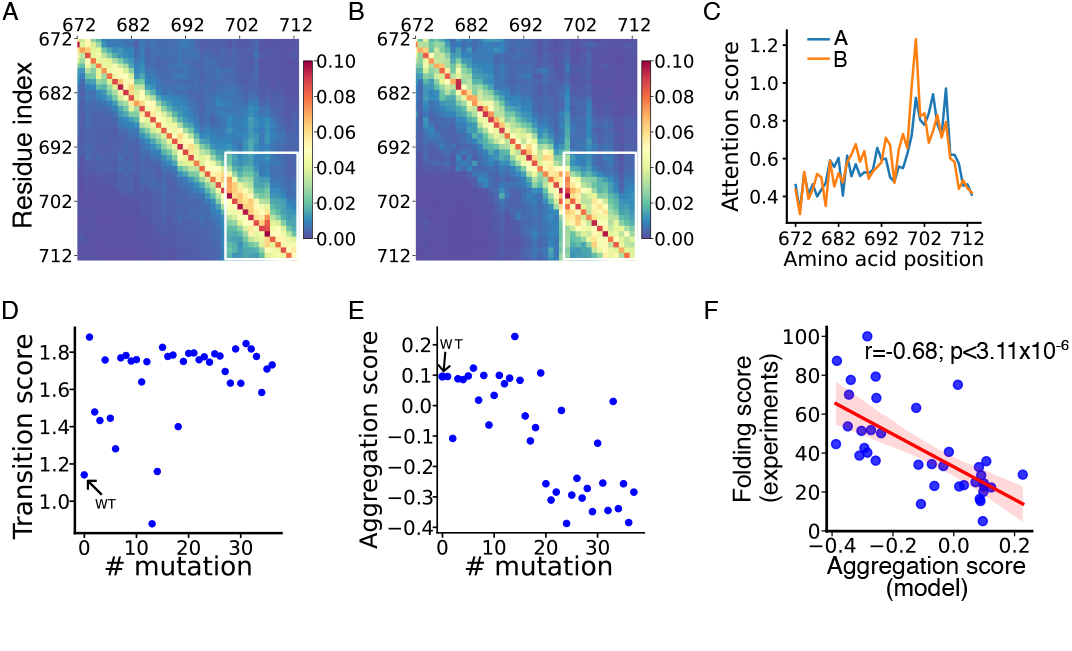
PPT predictions of *Aβ*42 variants. (A) and (B) Self-attention maps of classification tasks A and B reveal positions with a high density of *Aβ*42 variants. (marked by the white window). (C) Line plots show elevated attention between the 701-713 positions in APP for both tasks. (D) Predictions from classification task A indicate the tendency of *Aβ*42 variants to undergo PPT compared to the WT. (E) Predictions from classification task B show *Aβ*42 variants’ aggregation propensity compared to the WT. (F) Folding score versus predicted aggregation propensity for *Aβ*42 variants. Higher aggregation propensity is associated with a lower folding score, indicating a misfolded structure.

Overall, our modeling framework is useful for assessing protein variants. In particular, it might be useful for designing antibodies that show a reduced aggregation propensity. Similar analysis for additional sequences is presented in the SI Appendix, Fig. S13. We note that we use the same ESM-2 modeling architecture as described in ref. (38); however, in ref. (38) mutation effects are predicted from an unsupervised method that does not explicitly take into account PPT behavior. Our modeling approach addresses this gap by specifically considering alterations in PPT behavior.

### Assessing the relationship between PPT and gene expression for two AD-related proteins

Our predictions for two proteins known to be pathological in AD, APP and tau, revealed their propensity to undergo phase transitions (Fig. 10). Specifically, APP was predicted to have a high propensity to form amyloid aggregates along with a medium transition score (i.e., persisting in the droplet state to a medium extent). This aligns with the nature of this protein to form amyloid peptides upon proteolytic cleavage, such as *Aβ*42, which is considered very toxic and could potentially serve as an early biomarker for AD (39, 40). This indicates that the products of APP might have a lower free energy barrier for amyloid formation. In contrast, tau protein was predicted as a +Droplet driver with a lower transition score, suggesting a preference for the droplet state. This aligns with tau’s lack of a well-defined structure and its enrichment with hydrophilic amino acids, which can facilitate LLPS. This is in accordance with the observation that phase-separated tau droplets can both serve physiological functions (e.g., microtubule assembly) and act as an intermediate toward aggregation (41). Hence, LLPS might be considered a significant initiating factor in the onset of age-related diseases. External factors, such as phosphorylation—a potent regulator of tau (42–44)—can further contribute to the conversion of tau droplets into solid aggregates. Taken together, these predictions support the idea that amyloid beta plaques produced from APP can promote the formation of tau entanglements arising from aged droplets, which, in turn, damage neurons (45).

**Fig. 10.**
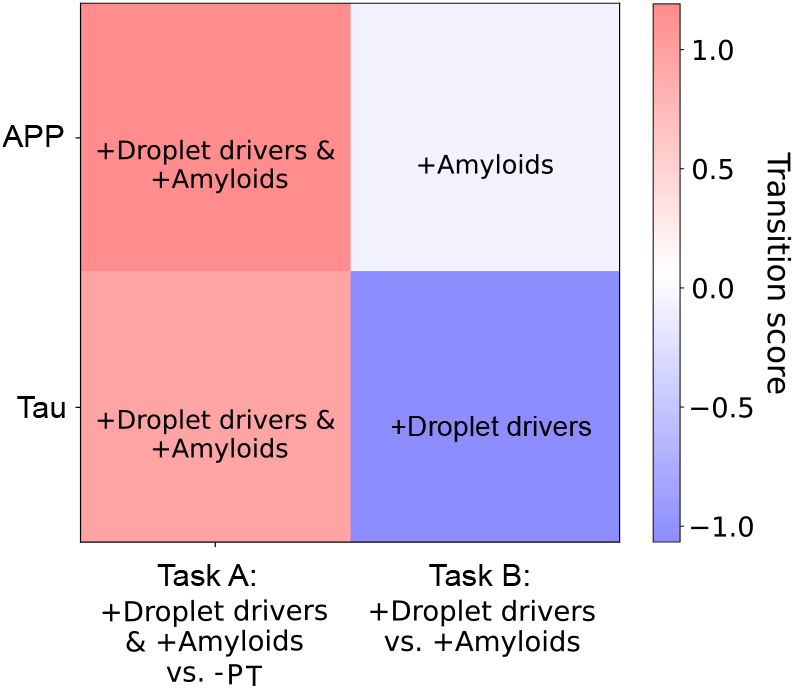
PPT prediction for APP and tau, two AD-related proteins. Transition scores, along with the predicted class, were calculated by the ML_LM model. Both proteins exhibited a strong propensity to undergo PPT (left panel). APP was predicted to have a high propensity to form amyloid aggregates, while tau was predicted to have a high propensity to form droplets (right panel).

We aimed to further understand how the propensity of tau and APP to form droplets and amyloid aggregates relates to gene expression and regulation (46). Our analyses revealed an overall decrease in gene expression in AD samples compared to controls (Fig. 11A). Additionally, we found that their transition scores correlate with the change in gene expression levels, indicating that a larger transition score is associated with a more significant decrease in gene expression (p<0.005 versus p<0.02). This suggests the existence of a negative feedback loop between protein levels and gene expression, tightly regulating protein aggregation in AD (and other neurodegenerative diseases), in accordance with (47, 48).

**Fig. 11.**
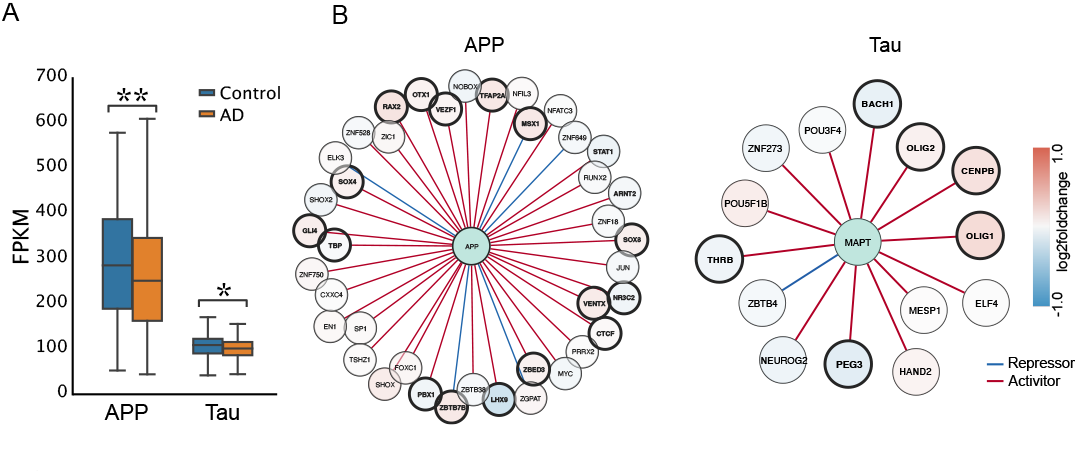
(A) Significant differences in gene expression were observed for both APP (left) and tau (right) proteins between AD samples and controls extracted from the dorsolateral prefrontal cortex. The age range of the samples spans from 67 to 90 years. ** p value < 0.005 and * p-value < 0.02. Significance was tested using the Welch statistical t-test. (B) Construction of the TF-gene regulatory network. In the center are APP and MAPT genes surrounded by their respective TFs. Blue TFs are downregulated while red TFs are upregulated in AD.

We further studied the feedback loop between gene expression and aggregation propensity by quantifying the regulatory mechanisms affecting APP and tau expressions. First, we conducted a differential gene expression analysis for the AD group and control, considering multiple covariates such as age, APOE genotype, and sex. We found that both APP and tau were downregulated in AD; however, APP showed a significant decrease (*p <* 3.8 *×* 10^−4^), while tau did not significantly change between the two groups. Second, we investigate the regulatory mechanisms of these genes by deriving the gene regulatory networks for APP and tau from DoRothEA (49), which include interactions between transcription factors (TFs) and target genes. As shown in Fig. 11B (dataset S9 and dataset S10), within these networks, APP or tau gene expression is regulated by two kinds of TFs: activators (nodes with red edges) and repressors (nodes with blue edges). We found that in the APP regulatory network, 45% of the TFs show significant differential expression (p<0.05). Among these TFs, all repressors are upregulated, and 37.5% of the activators are downregulated in the AD group. In the tau regulatory network, 43% of the tau TFs are significantly differentially expressed. In this case, only one repressor was found, which was not differentially expressed. Among the activators, 50% are downregulated, and the remaining 50% are upregulated. This suggests that proteins with a higher aggregation propensity are naturally subjected to more regulation to minimize their impact on the disease. In further support of this finding, we found evidence suggesting that APP could have a higher number of candidate cis-regulatory elements (cCREs) in the adult human brain than MAPT, the gene encoding for Tau (50). Specifically, when controlling for gene length, 0.24 cCREs were observed per kilobase pair that overlapped with APP, compared with 0.17 for MAPT. We note that this difference was primarily driven by an increase in candidate distal enhancers, which might influence the expression of other genes.

## Conclusions

To test the hypothesis that PPT is encoded by the amino acid sequence, we simulated the entire process from the soluble native state to the formation of droplets and amyloid aggregates. To this end, we fine-tuned an LLM and computed biophysical features, which were then used as input for classical machine learning models (e.g., RF classifier) to predict PPT. We highlighted the significance of self-attention-based language models in the prediction of PPT. Specifically, the LLM exhibited remarkably higher performance compared to the classical model benchmark. This emphasizes the capability of contextualized language models to decipher the functionality encoded by the amino acid sequence. In addition, we demonstrated that PPT is dictated by biophysical properties that can be directly computed from the protein sequence. In this regard, we showed that phase-separated proteins are less stable and more disordered. Furthermore, proteins associated with reduced solubility, high compactness, and increased hydrophobicity might be more prone to form amyloid aggregates. Our ML_LM model also showed high performance and proved to be useful in predicting sequence variants, suggesting that a combination of knowledge-based and contextualized models can serve as an effective predictor for PPT. While implementing our ML_LM model on unlabeled protein sequences and calculating their transition scores, we identified proteins potentially inclined to undergo a phase transition, forming droplets in particular. We also applied our modeling approach to tau and APP proteins. Both of these proteins are thought to contribute to AD, but there are different hypotheses as to which aggregates first. Our findings indicate that in terms of protein sequence, tau protein exhibits a higher propensity to form droplets compared to APP. This suggests that tau tangles might predominantly manifest in the advanced stages of AD, with APP plaques being more prevalent in the early stages of the disease. Moreover, we demonstrated that in AD, the phase transition propensity of these two proteins can be modulated by gene expression and TF activity levels. Specifically, we showed that APP is more prone to aggregation than tau and is commensurately down-regulated in AD. Taken together, our modeling approach can identify potential phase-separated proteins along with their variants, including those with a high propensity to be trapped in the liquid state, which are of great interest as they can serve as early biomarkers and drug targets (51) to slow the progression of age-related diseases. In this regard, time-dependent PPT experiments are still needed to leverage droplet formation as a stand-alone state and to discern its relationship with physiological and pathological conditions.

## Materials and Methods

### Database construction of +Droplet drivers and +Amyloids

The *+Droplet drivers* and *+Amyloids* datasets were constructed using previously published databases with no further modifications (15). These two databases contain proteins with UniProt accession identifiers, where each protein was labeled as a droplet driver or an amyloid based on the evidence to form droplets or amyloid aggregates under various experimental conditions.

**The droplet-forming protein** database was constructed from 404 droplet driver proteins that undergo spontaneous LLPS (without the aid of a partner) and are not reported to form amyloid aggregates. The database was assembled from three public databases: PhaSepDB (http:/db.phasep.pro) (52), PhaSePro (https://phasepro.elte.hu) (53), and the LLPSDB database (http://bio-comp.org.cn/llpsdb) (41). The material state that was experimentally observed for such proteins was liquid or gel. These proteins tend to have a high free energy barrier preventing them from transitioning to amyloid formation (12). We refer to these proteins as having a high propensity to form liquid condensates and label them as *+Droplet drivers* (dataset S1 and Fig. S1).

**The amyloid-forming protein** database was constructed from 113 proteins consisting of core amyloid regions. These amyloids can be either functional or pathological (http://amypro.net) (54). This dataset is enriched with experimentally validated amyloidogenic sequence regions. LLPS was not observed for these proteins, suggesting a low free energy barrier to rapidly form amyloid aggregates from liquid condensates, which cannot be readily observed. We refer to the proteins in this database as having a high propensity to form solid aggregates and label them as *+Amyloids* (dataset S2 and Fig. S2).

### Database construction of -PT

We randomly selected 360 sequences from the PDB* dataset constructed in (8). We intentionally did not use a larger number of sequences to maintain relative balance across the dataset. These sequences fold into a well-defined structure with high stability and are not enriched in any disordered residues, as indicated by the low fraction of disordered regions and instability index in Figs. 4A, 4C, dataset S3 and Fig. S3). The sequences were extracted from the PDB (55).

### Computing biophysical features from protein sequences

To examine whether biophysical properties are encoded in the amino acid sequence and can drive PPT, we calculated the following quantities to be fed into a classical machine learning model benchmark (e.g., RF model). First, we evaluated disordered regions using the IUPred3 algorithm, which predicts the probability for each amino acid in the sequence to be disordered using an energy estimation method (56). Next, we followed a similar process described in (8). Specifically, in each protein sequence, we identified regions with at least 20 consecutive amino acids with a probability above 0.5, considering these regions as disordered regions. Finally, we calculated the fraction of these regions. Second, to account for the 3D structure of the protein and its compactness in the 3D space, we calculated the gyration radius (29) as follows (20, 25):

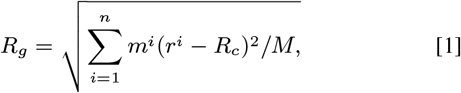

where *m*^*i*^ is the mass of the i-th atom, which is located in 3D space by the *r*^*i*^ coordinates, *Rc* is the coordinates of the center of mass, and *M* is the mass of all the atoms in the protein. This calculation was performed using PDB files, which contain information about each of the protein atoms’ coordinates in 3D space. The files were extracted from the AlphaFold database for the +Droplet drivers and the +Amyloids datasets, as their UniProt accession identifier was provided. If a PDB file was missing, it was constructed by ESMFold (v1) (20). In addition, we took advantage of the fast computing of protein structures by ESMFold and used this model to compute the PDB files of the -PT proteins, as their UniProt accession identifiers were not provided. We note that employing AlphaFold for predicting the 3D structure might result in an overestimation of the calculated gyration radius, especially for regions that are intrinsically disordered. However, we decided to consider the gyration radius as a feature because the predictions are biased in a uniform fashion. Furthermore, gyration radius is an important feature from a structural viewpoint and has proven to be useful in the predictions. Third, we calculated the instability index, *II*, a statistical measure of a given protein sequence, which is calculated by a weighted sum of dipeptides based on their differing occurrence in stable and unstable proteins as follows (26):

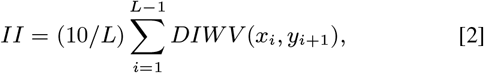

where *L* is the length of the protein sequence, *xi, yi*+1 is a dipeptide, and *DIWV* is the dipeptide instability weight value. A *II* below 40 is considered a stable protein. We next estimated the probability of solubility of all the protein sequences by using the solubility weighted index, *SWI*, calculated by taking the weighted average of the normalized B-factor of amino acid residues. Using the *SWI*, the probability of solubility is calculated as follows (27):

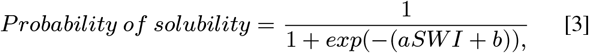

where *a* and *b* are fitting parameters. Finally, we calculated the mean hydrophobicity of each protein sequence using the Kyte and Doolittle hydropathy scale, where each amino acid is represented by a hydrophobicity score between -4.6 and 4.6. A score of 4.6 is the most hydrophobic, while a score of -4.6 is the most hydrophilic (28).

### ESM-2 - Transformer protein language model

We used the transfer learning approach by fine-tuning the attention-based ESM-2 model transformer with a linear layer on top for the sequence classification task (20). The model is composed of 6 layers and 8M parameters. We found this approach useful when dealing with relatively small datasets, as we have here. Specifically, we trained the model on our datasets using an 80:20 training-test set ratio. The model was trained with a batch size of 8 and 2 epochs. In this LLM, each amino acid is represented by a 320-dimensional embedding vector. We repeated this process separately for each of the two classification tasks described in Fig. 1.

### Random forest - Machine learning classifier

First, we normalized our features by performing min-max normalization. Next, we trained the RF model using the Python scikit-learn package (57), setting the class weight to ”balanced” and limiting the max depth of the trees to 5 to avoid overfitting. The model followed the same workflow as the LLM, as described in Fig. 1.

### Ensenble model: Random forest and LLM (ML_LM)

For each classification task, the combined probability of belonging in a certain class was calculated by averaging the probabilities of the RF model and the LLM. To label the sequences, we set a minimum threshold of 0.6 to be classified as -PT and +Droplet drivers for classification tasks A and B, respectively.

### Transition score calculation

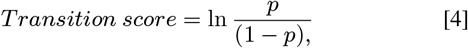

where *p* is the average probability of undergoing PPT, calculated by using the RF and LLM predictions.

We define the Δ transition score as the difference between the transition score of the mutant and the WT:

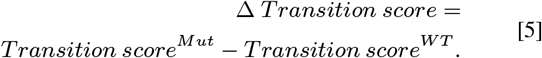

For task, A positive and negative Δ transition scores indicate higher and lower PPT propensity, respectively. For task B, a negative Δ transition score is associated with higher droplet formation (or lower aggregation), while a positive Δ transition score is linked to lower droplet formation (or higher aggregation).

### Locality parameter

We define the locality parameter *Lw* as the ratio between the median of the attention scores and the mean distance of the top five scores (*Si*) from the target amino acid within a sliding window size of 5×5 as:

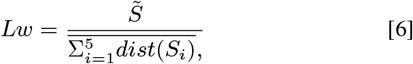

where 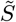 is the median attention score and *dist*(*Si*) is the distance from the target amino acid located on the diagonal.

### Benchmarking of mutated *Aβ*42 sequences

We focus on the *Aβ*42 peptide, which is relatively short compared to the full standard sequences used to train the models. Hence, we found it useful to use the ML_LM model to evaluate the aggregation propensity upon mutations. We note that because the LLM was trained on a broader context than 42 positions, we included the 50 amino acids before and after *Aβ*42 appear within the APP sequence. We also refined the ”fraction of disordered regions” feature to capture shorter subsequences within *Aβ*42 that are associated with disordered regions.

## Data availability

The data used or generated in this work is provided in the following files. The code is available from GitHub: https://github.com/gersteinlab/PPT.

**dataset S1**

**dataset S2**

**dataset S3**

**dataset S4**

**dataset S5**

**dataset S6**

**dataset S7**

**dataset S8**

**dataset S9**

**dataset S10**

## ACKNOWLEDGMENTS

We would like to thank Prof. Andrew Miranker from the Department of Molecular Biophysics and Biochemistry, Yale University, for fruitful discussions in relation to protein phase transitions from an experimental viewpoint. His expertise was helpful in shaping the direction of this work.

