## Supporting information for "Leveraging a large language model to predict protein phase transition: a physical, multiscale and interpretable approach"

#### This PDF file includes:

Supporting text
Figs. S1 to S13
Tables S1 to S2
Legends for Dataset S1 to S10
SI References

#### Other supporting materials for this manuscript include the following:

Datasets S1 to S10

### Supporting Information Text

#### ESMFold - Transformer protein language model

We fine-tuned the ESM-2 (1, 2), a large language model trained on a masked language modeling objective. We set the maximum length of the tokenizers to 1,100 amino acids and padded the rest of the protein sequences with zeros up to their maximum length. This setting allowed us to run the model using 3 GPUs across 3 nodes (1 GPU per node) and enabled us to fully tokenize almost all of our sequences.

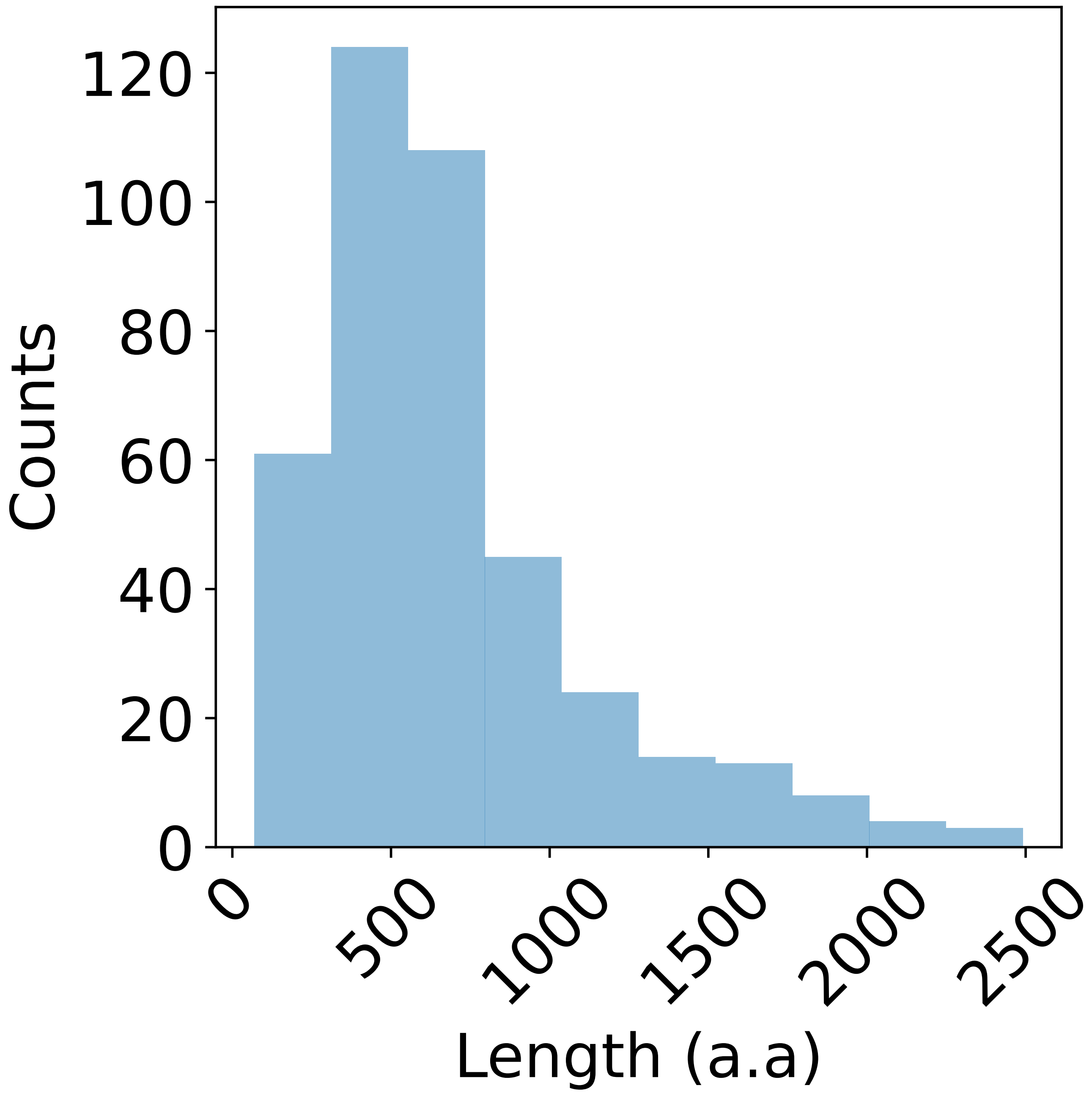


**Fig. S1.** Histogram of amino acid lengths. Class: +Droplet drivers.

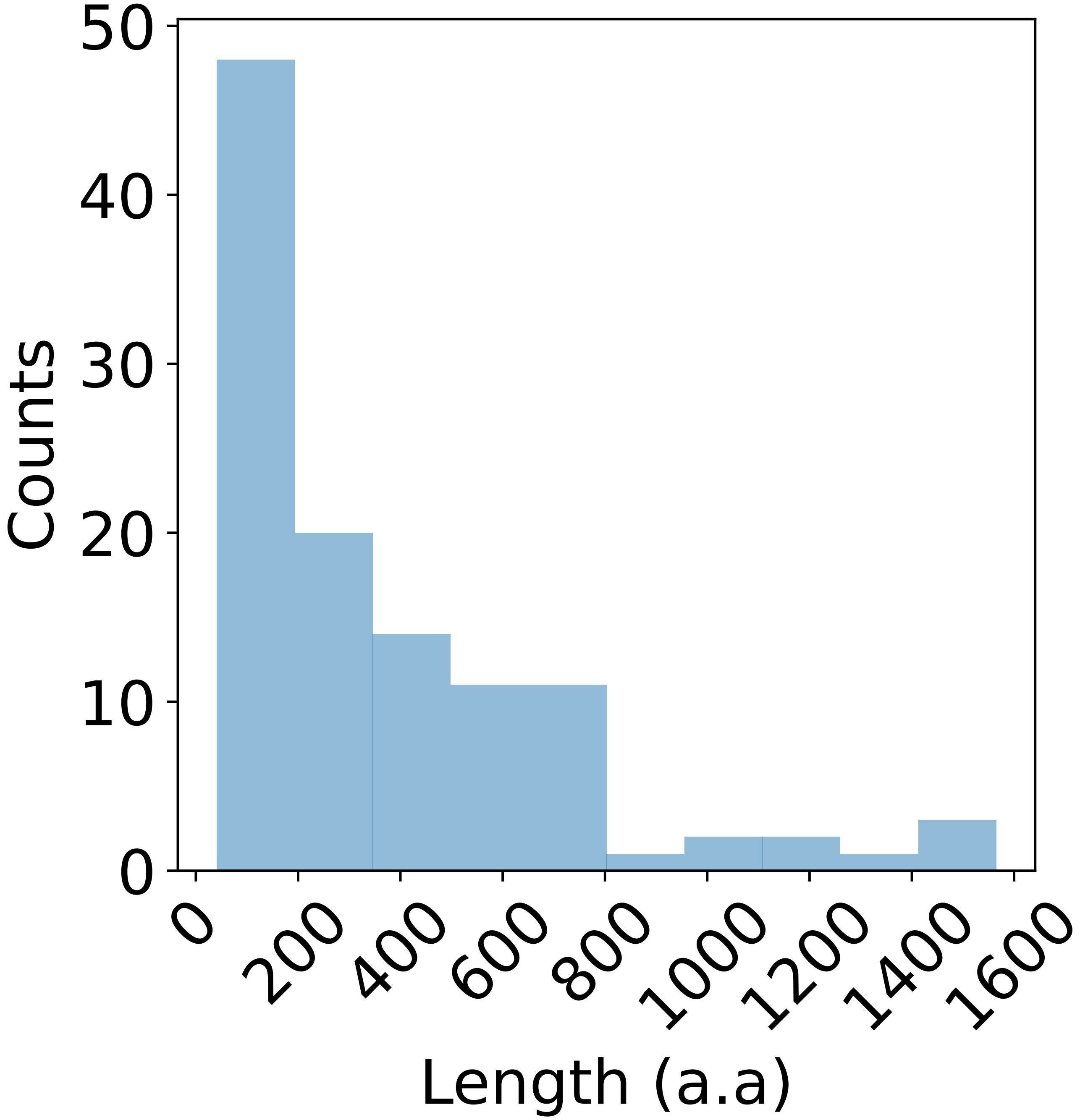


**Fig. S2.** Histogram of amino acid lengths. Class: +Amyloids.

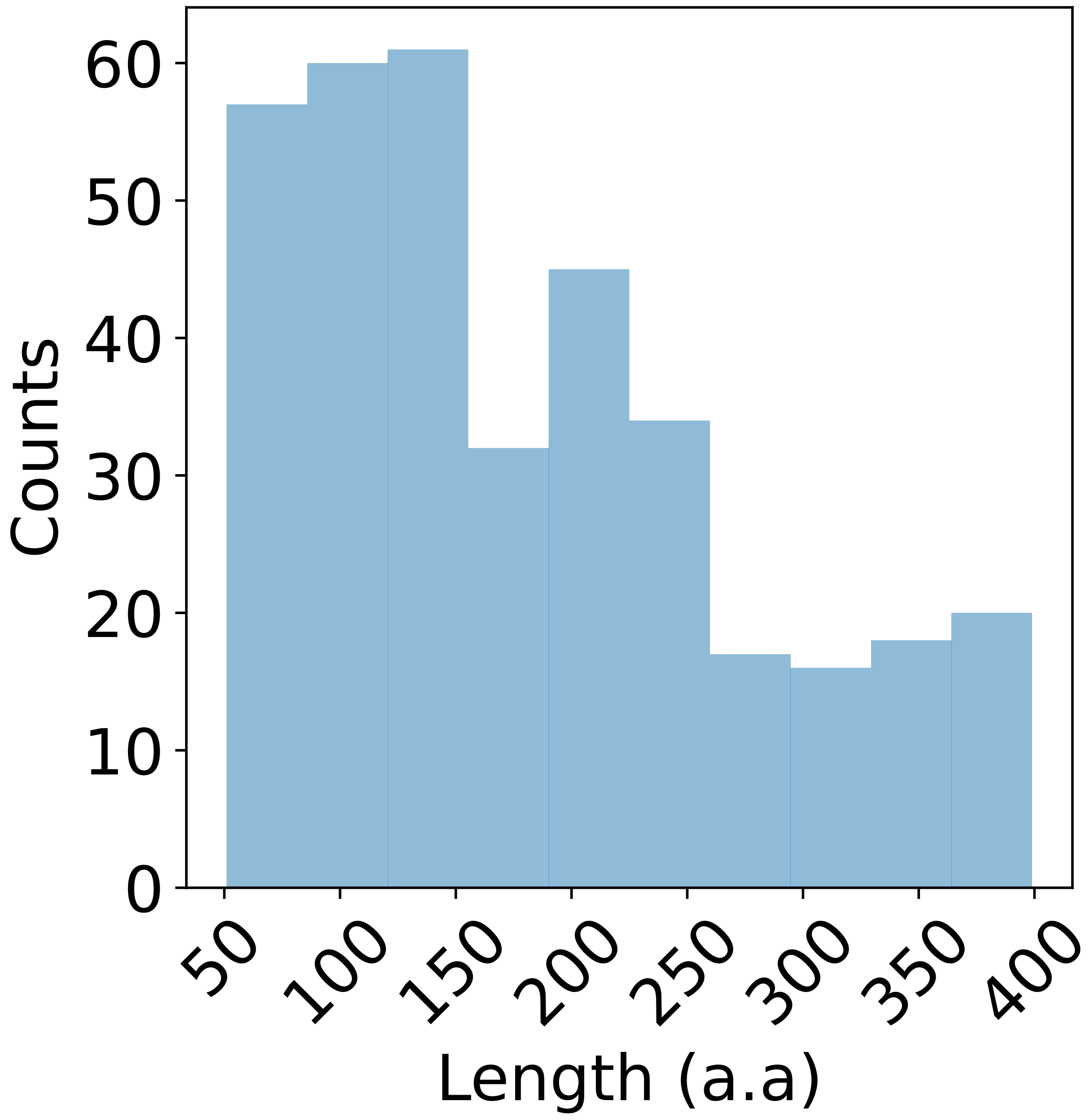


**Fig. S3.** Histogram of amino acid lengths. Class: -PT.

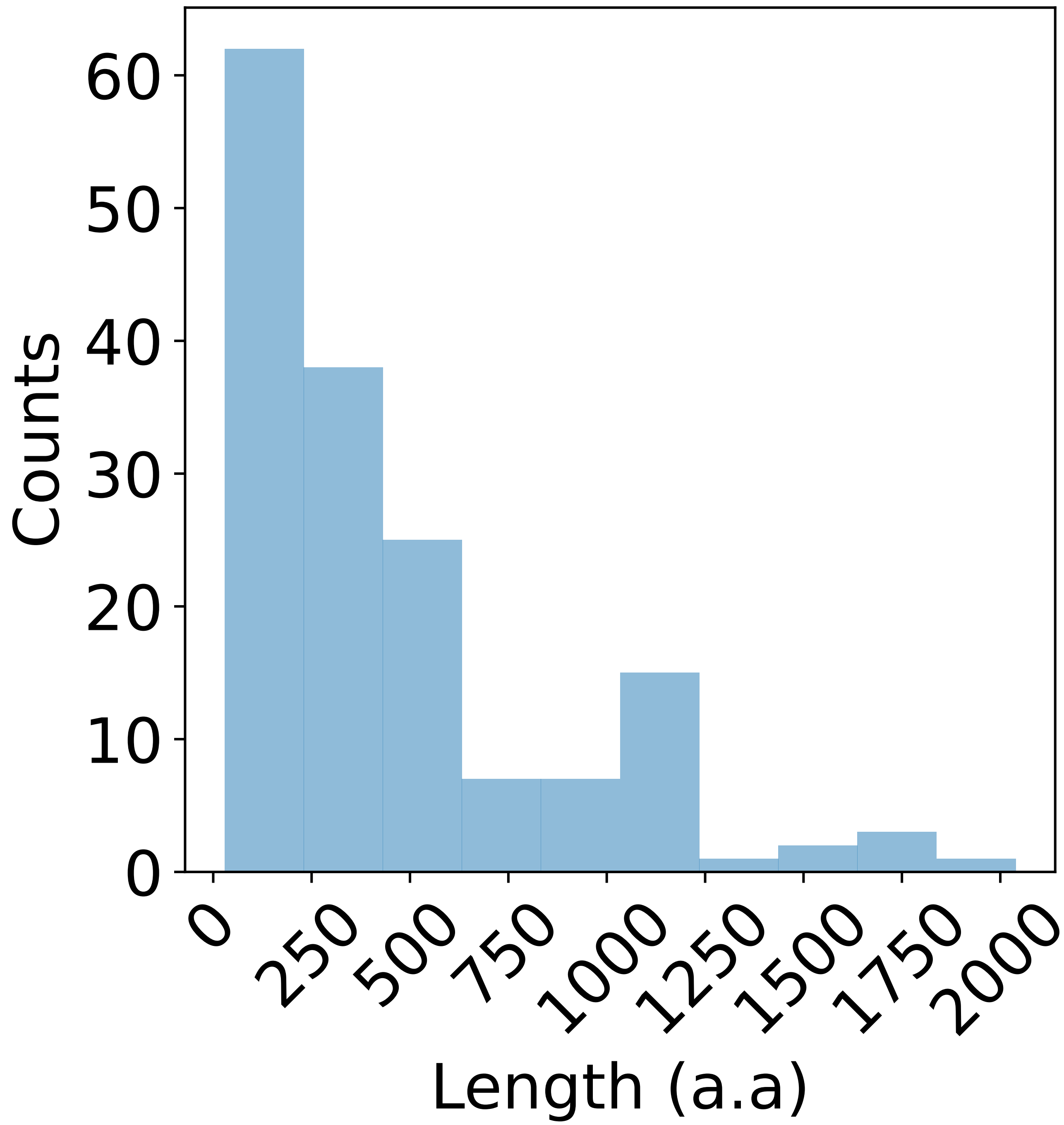


**Fig. S4.** Histogram of amino acid lengths. Class: Unlabeled sequences.

**Table S1. Classification Task A: Five-fold cross-validation for classical model benchmark.**

| Model | Accuracy | Recall | Precision | AUROC |
|---|---|---|---|---|
| 1. Logistic regression | 0.85±0.033 | 0.85±0.03 | 0.85±0.031 | 0.92±0.028 |
| 2. SVM (radial basis function) | 0.87±0.038 | 0.87±0.035 | 0.86±0.037 | 0.94±0.024 |
| 3. SVM (linear) | 0.83±0.044 | 0.84±0.041 | 0.83±0.043 | 0.92±0.033 |
| 4. Gaussian Naive Bayes | 0.77±0.052 | 0.79±0.05 | 0.79±0.046 | 0.84±0.07 |
| 5. Random forest | 0.86±0.046 | 0.86±0.046 | 0.86±0.047 | 0.94±0.027 |
| 6. Large language model | 0.96±0.021 | 0.96±0.020 | 0.95±0.022 | 0.96±0.02 |

**Table S2. Classification Task B: Five-fold cross-validation for classical model benchmark.**

| Model | Accuracy | Recall | Precision | AUROC |
|---|---|---|---|---|
| 1. Logistic regression | 0.73±0.048 | 0.71±0.026 | 0.66±0.025 | 0.79±0.05 |
| 2. SVM (radial basis function) | 0.8±0.033 | 0.79±0.037 | 0.72±0.035 | 0.88±0.021 |
| 3. SVM (linear) | 0.75±0.048 | 0.74±0.021 | 0.69±0.027 | 0.8±0.038 |
| 4. Gaussian Naive Bayes | 0.83±0.028 | 0.7±0.073 | 0.75±0.041 | 0.86±0.037 |
| 5. Random forest | 0.84±0.027 | 0.75±0.054 | 0.77±0.038 | 0.88±0.02 |
| 6. Large language model | 0.89±0.03 | 0.77±0.053 | 0.89±0.051 | 0.93±0.036 |

**SI Dataset S1 (datasetS01.csv)**
Proteins prone to droplet-driver formation, labeled as +Droplet drivers (3).

**SI Dataset S2 (datasetS02.csv)**
Proteins prone to amyloid formation, labeled as +Amyloids (3).

**SI Dataset S3 (datasetS03.csv)**
Proteins unlikely to undergo phase transitions, labeled as -PT (4).

**SI Dataset S4 (datasetS04.csv)**
Unlabeled protein sequences (5).

**SI Dataset S5 (datasetS05.xlsx)**
Classification Task A: PPT predictions for unlabeled protein sequences.

**SI Dataset S6 (datasetS06.xlsx)**
Classification Task B: PPT predictions for unlabeled protein sequences.

**SI Dataset S7 (datasetS07.csv)**
Classification Task A: Predicted transition scores for $A\beta42$ variants.

**SI Dataset S8 (datasetS08.csv)**
Classification Task B: Predicted transition scores for $A\beta42$ variants.

**SI Dataset S9 (datasetS09.csv)**
APP regulatory network.

**SI Dataset S10 (datasetS10.csv)**
Tau regulatory network.

### 1. Attention maps for point mutations

Given the high sensitivity of large language models to local changes in the sequence (e.g., point mutations) compared to classical ML approaches (when dealing with moderate-to-long sequences), we used the self-attention maps to analyze three FUS ALS-associated point mutations (P525L, R521C, and R244C) that were shown experimentally to enhance droplet formation (6). A local elevated attention in the amino acid mutated position or nearby is observed, implying a change in the local interaction patterns that can drive or hinder phase transition (S5-S7:A and C, S5-S7:B and D). To demonstrate the local change in the attention score upon mutation, we computed the absolute difference between the attention maps of the mutated sequence and the wild type (WT), followed by summation across the columns to generate a line plot over the residue positions, as presented in Figs. S5-S7: E and F.

(1) P525L:

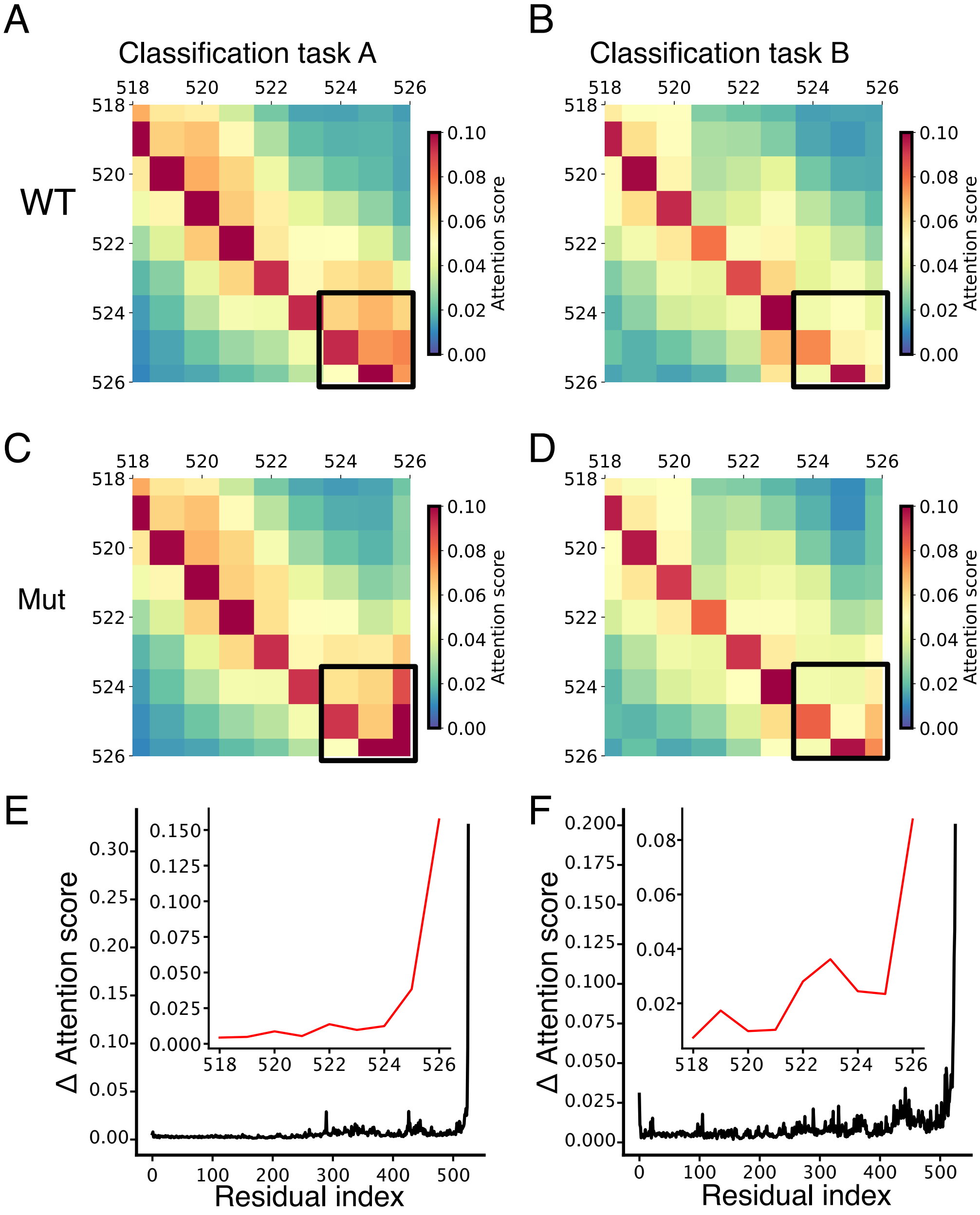


**Fig. S5.** FUS attention maps extracted from classification tasks A and B for the WT versus the P525L point mutation. (A) and (B) show the attention maps for the WT. (C) and (D) show the attention maps for the P525L mutant. Local attention around the mutation position is marked by the black rectangle. (E) and (F) are line plots showing elevated attention upon mutation relative to the WT for classification tasks A and B, respectively.

 

(2) R521C:

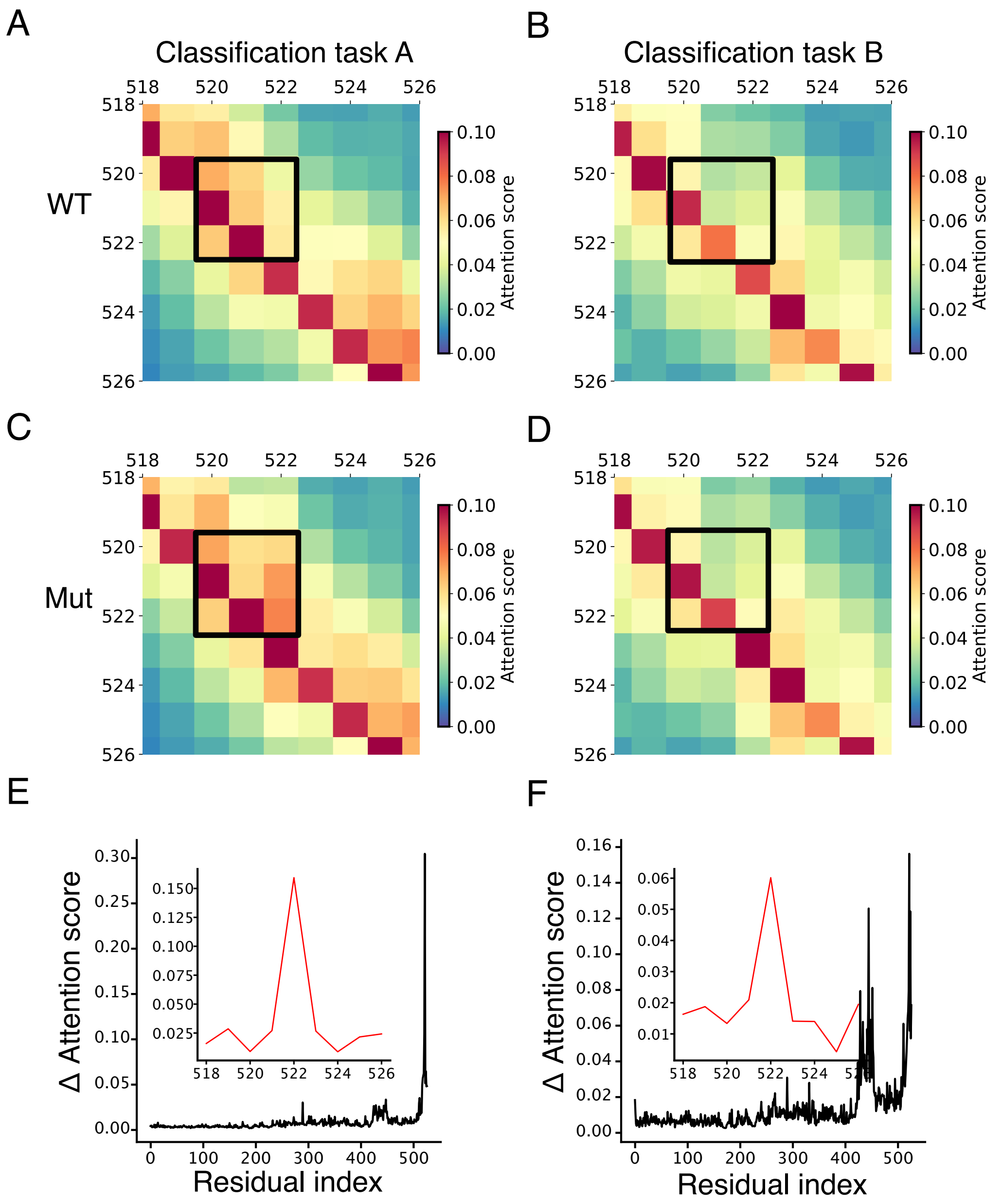


**Fig. S6.** FUS attention maps extracted from classification tasks A and B for the WT versus the R521C point mutation. (A) and (B) show the attention maps for the WT. (C) and (D) show the attention maps for the R521C mutant. Local attention around the mutation position is marked by the black rectangle. (E) and (F) are line plots showing elevated attention upon mutation relative to the WT for classification tasks A and B, respectively.

(3) R244C:

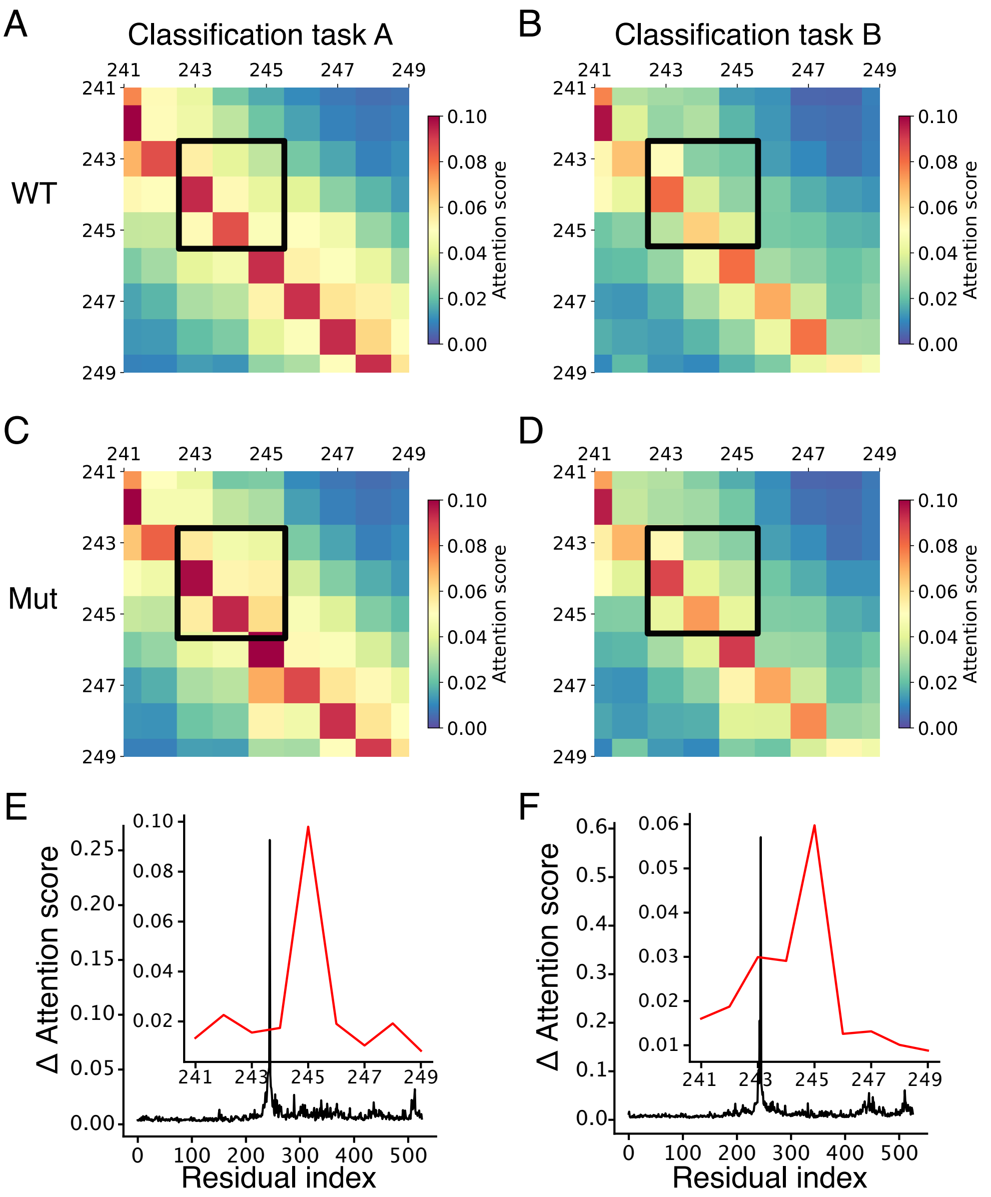


**Fig. S7.** FUS attention maps extracted from classification tasks A and B for the WT versus the R244C point mutation. (A) and (B) show the attention maps for the WT. (C) and (D) show the attention maps for the R244C mutant. Local attention around the mutation position is marked by the black rectangle. (E) and (F) are line plots showing elevated attention upon mutation relative to the WT for classification tasks A and B, respectively.

### 2. Computing the Δ transition score between the mutant and WT

We utilize Eq. 5 to compute the difference between the transition score of the mutant and the WT, refered to as the Δ transition score:

(2.1) FUS (P35637) point mutations

We show that all three point mutations are more likely to undergo phase transition upon mutation (classification task A). Furthermore, all three mutants had a reduced aggregation score in classification task B, associated with a higher propensity to form more droplets compared to the WT (Fig. S8A). Our predictions align with the experimental measurements presented in (6), where more droplets were observed in P525L, R521C, and R244C than in the WT. The last mutant (R244C) was experimentally validated to enhance aggregation propensity, which is in accordance with its highest Δ transition score in classification tasks A and B, implying that this mutant possesses the ability to gradually aggregate within the droplets.

(2.2) FUS 1-214 IDR region

We further investigated the FUS 1-214 intrinsically disordered region (IDR) by applying mutations reported in (7), where tyrosine is switched with serine: 5Y->S, 9Y->S, and 27Y->S, as shown in Fig. S8B. The propensity for phase transition was maintained but diminished upon changing from tyrosine (Y) to serine (S), with the most significant reduction observed in the 27Y to S substitution (classification task A). The propensity for droplet formation (classification task B) decreased with a higher frequency of switching from tyrosine (Y) to serine (S), as demonstrated by the heatmap. These results align with (7), highlighting the importance of aromatic residues, particularly tyrosine, in IDR-mediated phase separation.

(2.3) hnRNPA1 (P09651) PLCD region

We performed predictions for the prion-like low complexity domains (PLCD) region as described in (8). As shown in the heatmap below (Fig. S8C), in classification task A, we observed a minor decrease in phase transition propensity upon the Tyr (Y) -> Phe (F) mutation and vice versa. The propensity for the (Y)->(F) mutation to undergo a phase transition was reduced further. Substituting (F) with (Y) residues enhanced the propensity for droplet formation, while replacing (Y) with (F) showed a reduced ability to form droplets, which is associated with a higher transition score in classification task B. Our predictions align with (8), where substituting (F) with (Y) residues enhanced the driving forces for phase separation, but the opposite weakened the phase separation propensity.

(2.4) G3BP1 (Q13283)

Substituting the five arginines (R) in the RGG region with phenylalanine (F) disrupts droplet formation, as evidenced by a higher Δ transition score in classification task B (Fig. S8D). Our prediction aligns with (9), where G3BP1 containing (R) to (F) mutations at all five arginine sites was defective in cytoplasmic condensate formation under untreated conditions.

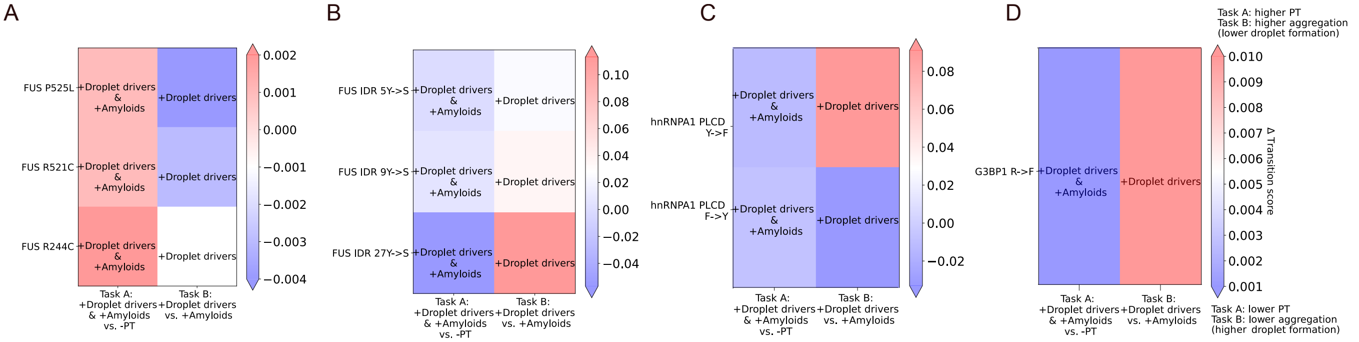


**Fig. S8.** Changes in the transition score upon mutations. (A) FUS ALS-related point mutations. Higher droplet formation is predicted upon mutation. (B) Mutants within the FUS 1-214 IDR region. The propensity to undergo phase transition is predicted to be reduced for all the three mutations. Higher mutation frequency correlates with a greater reduction in phase transition propensity. (C) Replacing F with Y is expected to increase the propensity for phase transition, resulting in the formation of more droplets. (D) Substituting the five arginines (R) in the RGG region with phenylalanine (F) disrupts droplet formation. PT is an abbreviation for phase transition propensity.

### 3. Protein design based on phase transition propensity

Based on expert knowledge, regions in protein seqeunces can be excluded by eliminating attention (i.e, masking) of multiple amino acid positions suspected to contribute to phase transition behavior. By masking different amino acid positions in the large language model and observing the model's predictions, experimentalists can gain insights into how the model interprets different parts of the sequence and their importance for phase transitions. Use cases are provided by masking regions in FUS and amyloid beta precursor (APP) proteins that were validated to substantially contribute to the propensity for phase transition. Specifically, by computing the Δ transition score, our predictions show that masking the IDR region in the FUS protein (7) diminishes the propensity for droplet formation (Fig. S9A). Similarly, by masking known regions (a region containing the $A\beta42$ peptide and the C-terminal) in APP, we predict a diminished propensity to form droplets and amyloids, respectively (Figs. S9B and C), which is in line with (10–12). Our code implements the masking option through a function called "masking_regions."

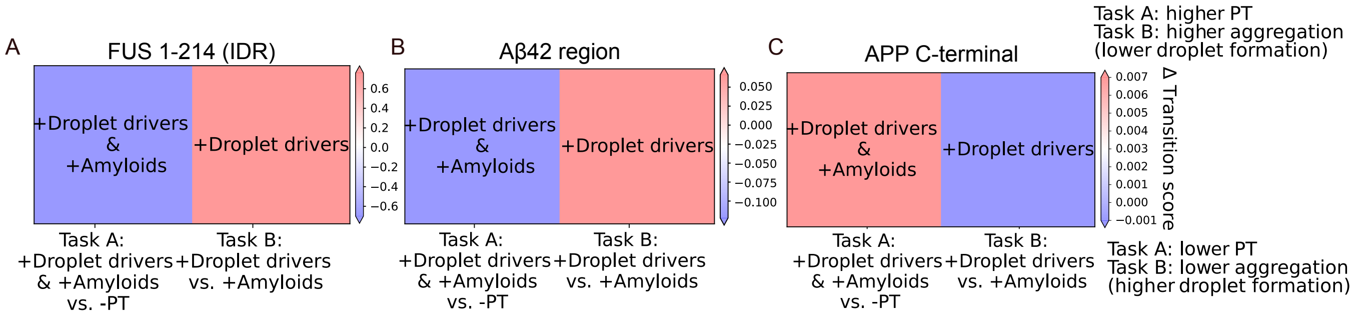


**Fig. S9.** Masking regions in FUS and APP sequences. (A) Masking the intrinsically disordered region (1-214) in FUS results in reduced droplet formation. (B) Masking the $\alpha\beta42$ region in APP leads to reduced droplet formation. (C) Masking the C-terminal region of APP reduces the tendency of APP to form amyloids while enhancing the propensity for droplet formation.

### 4. Quantification of local attention versus sparse attention

We applied the locality parameter (Lw) to FUS (P35637), hnRNPA1 (P09651), and G3BP1 (Q13283). In classification task A, we consistently observed local attention, while in classification task B, sparse attention was evident, corresponding to high and low Lw values (p-value<0.05, Fig. S10).

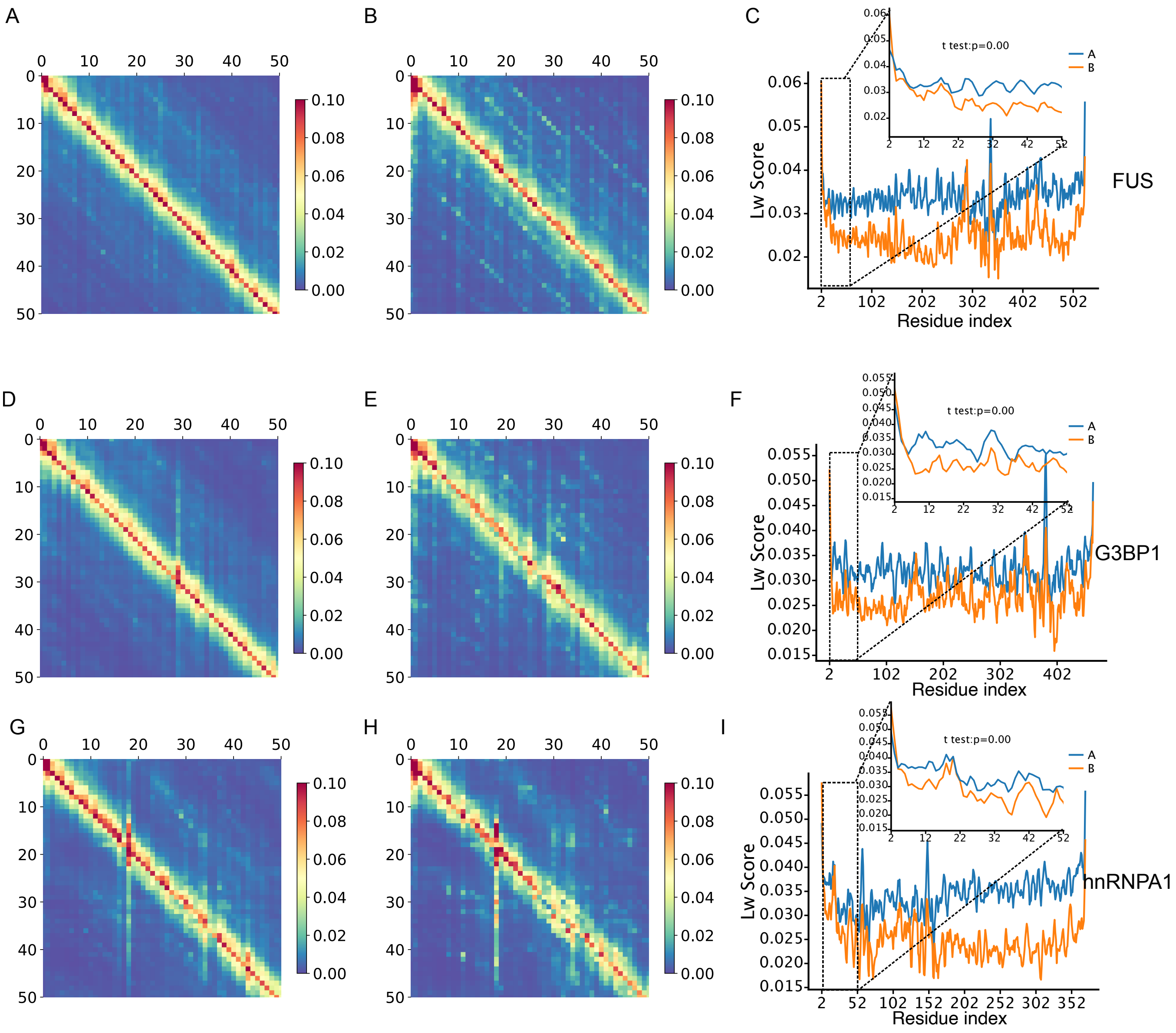


**Fig. S10.** Attention maps for FUS, G3BP1, and hnRNPA1 for classification tasks A and B (left and middle panels, respectively). In all three proteins, local attention is evident in classification task A, whereas sparse attention is observed in classification task B. This distinction is quantified by the Lw line plots displayed in the right panel. (A), (B), and (C) refer to FUS. (D), (E), and (F) refer to G3BP1. (G), (H) and (I) refer to hnRNPA1.

### 5. Learning criteria by the large language model

In classification task A, the embedding vectors of +Droplet drivers and the +Amyloids classes partially overlap (Fig. S11). This implies that classification task B incorporates additional learning criteria beyond those that have already been utilized in classification task A to maximize the difference between the two classes.

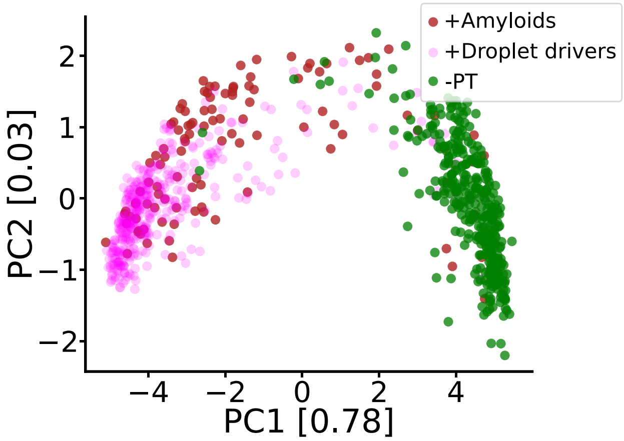


**Fig. S11.** PCA of the embedding vectors of classification task A. A partial overlap of the data points is observed between the +Amyloids and the +Droplet drivers classes.


Mor Frank, Pengyu Ni, Matthew Jensen and Mark B Gerstein

### 6. Statistical considerations for feature selection

We used features that are commonly employed in the field of protein phase transition. Specifically, we chose to concentrate on a limited set of biophysical, structural, and thermodynamic features that are easily interpretable, in contrast to a large language model, like ESMFold, which inherently analyzes the encoded grammar. We note that features lacking significance between +Droplet drivers and +Amyloids were excluded from the analysis, considering their relatively minor relevance in distinguishing between these two classes (Fig. S12).

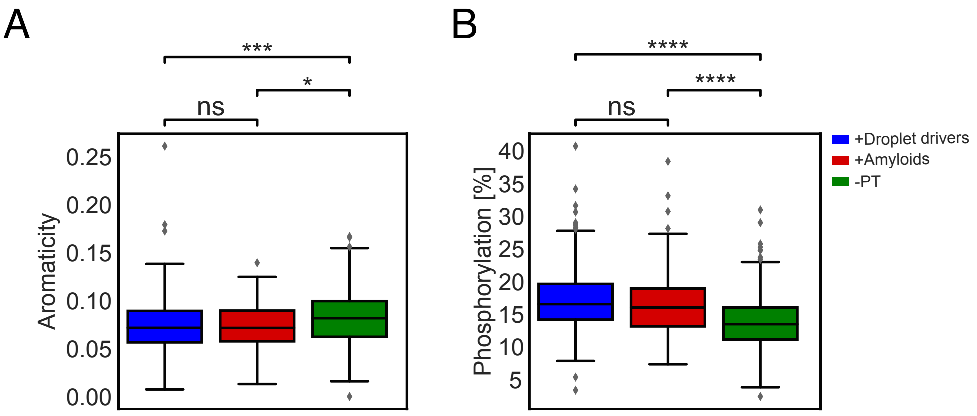


**Fig. S12.** Representative examples of non-significant features (ns). (A) Aromaticity: relative frequency of Phe, Trp and Tyr. (B) Phosphorylation: percentage of serine, threonine, and tyrosine. The distinction between +Droplet drivers and +Amyloids is not significant.

### 7. Predictions of unlabeled protein sequences

We applied our modeling framework to 189 unlabeled standard protein sequences extracted from the AMP-AD database (5) (dataset S4), with predictions derived from the ML_LM model. The transition score of +Droplet drivers and +Amyloids versus -PT revealed two distributions. Most of the protein sequences were predicted to have a high propensity to undergo a phase transition, forming either droplets or amyloid aggregates, which is in line with previous findings (13) (Figs. S13A and S13C, and dataset dataset S5). Next, we further evaluated the propensity of these proteins to form either +Droplet drivers or +Amyloids. Our analyses demonstrated that most proteins exhibited a high propensity to undergo a phase transition, mainly forming droplets (Figs. S13B and S13D, and dataset dataset S6), emphasizing the physiological relevance of droplets (14–16).

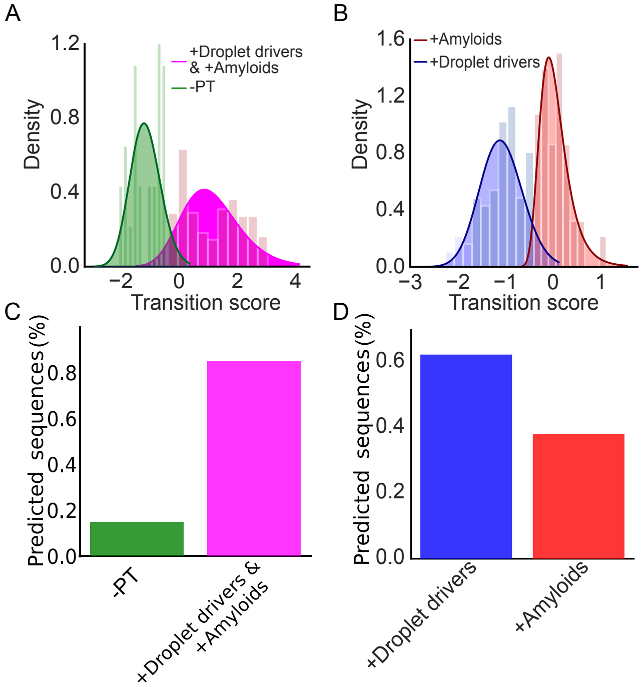


**Fig. S13.** Probability density of the transition scores along with the fitted alpha distributions (solid line) calculated for the unlabeled protein sequences along the percentage of the predicted sequences for each classification task. (A) +Droplet drivers and +Amyloids are shifted to positive scores compared to -PT, which is concentrated in the negative range. (B) +Droplet drivers and +Amyloids are mostly located in the negative and positive ranges, respectively. (C) The majority of the unlabeled proteins were predicted to be +Droplet drivers and +Amyloids. (D) Most of the unlabeled protein sequences were predicted to have a high propensity to form droplets.
